# Pre-exposure to time-restricted feeding reduces tissue CD4^+^ T cells with a limited effect on *Mycobacterium tuberculosis* clearance at the early time point

**DOI:** 10.1101/2025.04.18.649546

**Authors:** Ashish Gupta, Nidhi Yadav, Subhasmita Das, R. Rajendra Kumar Reddy, Nupur Sharma, Amol Ratnakar Suryawanshi, Jaswinder Singh Maras, Ranjan Kumar Nanda

**Affiliations:** Translational Health Group, Internationalw Centre for Genetic Engineering and Biotechnology, New Delhi-110067, India; Clinical Proteomics Laboratory, BRIC-Institute of Life Sciences, Bhubaneswar-751023, India; ILS Mass Spectrometry Facility, BRIC-Institute of Life Sciences, Bhubaneswar-751023, India; Department of Molecular and Cellular Medicine, Institute of Liver and Biliary Sciences, New Delhi-110070, India

**Keywords:** Time-restricted feeding, *Mycobacterium tuberculosis*, metabolomics, immunology

## Abstract

Time-restricted feeding (TRF) in human and animal models positively impacts metabolic health by improving glucose homeostasis and reducing adiposity. However, limited studies have investigated the impact of TRF on the host immune response and upon bacterial infection. In this study, C57BL/6 mice (6–8 weeks old, male) were subjected to 8 hours of dark-phase TRF for 30 days, and aerosol infected with a low dose (100-400 colony-forming units) of *Mycobacterium tuberculosis* (Mtb) H37Rv. The body weight gain, multi-tissue metabolome, and proteome changes and distribution of immune cells in these study groups were monitored. In the first 15 days of TRF, mice showed better glucose tolerance with marginal weight loss. The serum and liver metabolome of TRF mice demonstrated perturbed fatty acid biosynthesis and degradation, steroid hormone biosynthesis pathways and tyrosine metabolism. Liver proteome data of TRF mice indicated an increased fatty acid oxidation which might affect the host immune cell frequency and functionality. The Mtb-infected TRF and control mice had similar tissue (lung, spleen and liver) bacterial burden at 21 days post-infection. The bone marrow of Mtb-infected TRF mice had significantly lower CD3^+^ T cells, and CD4^+^ T cells were low both in the bone marrow and lungs. This study reports that mice undergoing 8 hours of TRF had an improved metabolic phenotype, and discontinuing TRF limits its positive impact on handling Mtb infection at an early point.

## Introduction

Time-restricted eating or feeding (TRE/F) is a dietary intervention that restricts the time window of food intake to 6–12 hours a day in humans or animal models. This intervention increases the daily window of fasting, reducing their body weight and improving metabolic status in overweight, obese and prediabetic individuals.^1,2^ However, limited studies have investigated the impact of TRF on the immune system and whether it resolves or worsens infectious disease conditions like tuberculosis (TB) is unknown.

TRF improves the host metabolic parameters by synchronizing food availability with the expression of enzymes involved in nutrient utilization.^2^ Night-restricted TRF in mice, which limits food access to the lights-off phase, is reported to reduce fat mass, dampen high-fat diet-induced weight increase, and improve glucose homeostasis.^2,3,4,5^ TRF in mice is also reported to reduce inflammation by lowering the expression of inflammatory cytokines and chemokines, but limited reports have investigated its impact in the context of infection.^2,6–10^ In this study, we aimed to investigate the effect of 30 days of TRF on the immune system of male C57BL/6 mice and its impact on *Mycobacterium tuberculosis* (Mtb) H37Rv infection. TRF improves host metabolism at the tissue or cellular level, and establishing its role in resolving infections may help us to develop appropriate dietary interventions that benefit TB patients.

## Experimental Procedures

### Study Animals

In this study, all the experiments were performed following the approved procedures of the Institutional Animal Ethics Committee of the International Centre for Genetic Engineering and Biotechnology, New Delhi (ICGEB/IAEC/16072024/41.8). Male C57BL/6 mice (six to eight weeks old) were used in this study. Five animals per cage were housed and kept under a 12-hour light/dark cycle with lights on at 8 AM and off at 8 PM. Mice groups were randomly assigned to either ad libitum (ALF) or a time-restricted feeding (TRF) group. Mice in the TRF group had access to the food only for 8 hours from Zeitgeber time (ZT)13–ZT21, where ZT0 is the time when the lights are switched on and ZT12 lights were switched off. During the entire experimental duration, food intake and body weight of the mice were measured daily and weekly, respectively.

### Intraperitoneal glucose tolerance test (iPGTT)

After fasting the mice for 16 hours (ZT21 to ZT13), glucose (1 g/kg body weight) was introduced intraperitoneally and blood glucose levels were measured from the tail vein at different time intervals (0, 15, 30, 60, 90 and 120 min) using a glucometer (Dr. Morepen BG-03 Gluco One Glucometer). At the indicated time points, whole blood samples were collected via retro-orbital bleeding from the mice (after 16 hours of fasting) and, after 20 minutes, were centrifuged at 3,500 g for 20 min at 4°C to separate the serum, and then aliquoted and stored at -80°C for further analysis.

### Serum insulin and free fatty acid measurement

Fasting insulin levels were measured in the serum of mice after 16 hours of fasting, using the Rat/Mouse insulin ELISA kit (Millipore Sigma, Cat no. EZRMI-13K) following the manufacturer’s instructions. And homeostatic model assessment of insulin resistance (HOMA-IR) was calculated using the following formula:^11^

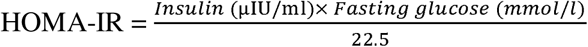

Fasting-free fatty acid levels in the serum of experimental mouse groups were measured using a fluorometric assay (Cayman Chemical, Cat no. 700310) following the manufacturer’s instructions.

### Serum and liver tissue metabolite isolation and mass spectrometry analysis

To the serum samples (50 µl), chilled methanol (400 µl) and internal standard (ribitol, 2 µl, 0.5 mg/ml) were added, and after vortexing, the samples were incubated at -20°C overnight. These processed serum samples were centrifuged at 16,000 g for 10 min at 4°C, and the supernatant was dried in a SpeedVac (Labconco). The dried pellet was resuspended in 5% acetonitrile and transferred to an autosampler vial for global metabolite profiling using liquid chromatography and mass spectrometry (LC-MS). The LC-MS data were acquired using a Thermo Scientific^TM^ UHPLC system combined with a Q Exactive Orbitrap Mass Spectrometer. Hypersil GOLD^TM^ C18 Selectivity HPLC Column was used for metabolite separation. The mobile phases consisted of 1% formic acid as solvent A and 100% acetonitrile. The gradient was 1 min, 5% solvent B; 17 min, 99% solvent B; 21 min, 99% solvent B; 22 min, 5% solvent B; 25 min, 5% solvent B. The solvent flow rate was set at 0.5 ml/min, and the column temperature was set at 60°C. The ion source temperature ranged from 250 to 300 °C, with a spray voltage of 5500–4500 V. The MS data were acquired in both positive and negative modes with a mass range of 150 to 2,000 Da, using nitrogen as collision gas, and at a resolution of 70,000 m/Δm. The LC-MS data were preprocessed using Compound Discoverer 3.3 to identify the metabolites.

Similarly, to the liver (50 mg) tissue, chilled methanol (400 µl, 80%), internal standard (ribitol, 2 µl, 0.5 mg/ml) and zirconium beads (2 mm, 250 mg) were added and homogenized in three cycles (30 sec on; 30 sec off) using a bead beater (Biospec Mini-Beadbeater-16). After bead beating, the extracted liver metabolites were incubated on ice for 30 min. Following centrifugation at 16,000 g for 10 min at 4°C, the supernatant was dried using a SpeedVac (Labconco). To these dried processed samples, methoxamine hydrochloride (20 mg/ml, 40 μl) was added and incubated at 60°C for 2 hours at 900 rpm in a thermomixer. To the reaction mixture, N-methyl-N-(trimethylsilyl)trifluoroacetamide (MSTFA, 70 μl) was added and incubated at 60°C for 30 minutes at 900 rpm. After incubation, the samples were centrifuged at 10,000 g for 10 minutes at 25°C, and the supernatant was transferred to gas chromatography (GC) vial inserts for data acquisition using a 7890 Gas Chromatograph coupled to a Pegasus 4D GC×GC-time-of-flight mass spectrometer. The derivatized samples (1 μl) were injected into an HP-5 ms column (30 m length, 0.25 mm width, 250 μm internal diameter) in splitless mode using helium as a carrier gas at a constant flow rate (1 ml/min). The secondary column was Rxi-17 (1.5 m length and 250 μm diameter). Electron ionization mode was fixed at −70 eV to scan ions of 33 to 600 m/z range at an acquisition rate of 20 spectra/second. The ion source temperature was set at 220°C. The GC oven parameters used for acquisition were as follows: 50°C hold for 1 min, temperature increased to 200°C with a ramp of 8.5°C/min, further increased to 280°C with a ramp of 6°C/min and a hold time for 5 min. The secondary oven temperature offset was set at 5°C relative to the GC oven temperature, and the modulator temperature offset was set at 15°C relative to the secondary oven temperature. The transfer line temperature was set at 225°C. A solvent delay of 600 s was used during data acquisition, and GC-MS data of derivatized samples were acquired within 24 h of derivatization. All GC-MS raw data files of the study groups were aligned using the “Statistical Compare” feature of ChromaTOF (4.50.8.0, Leco), and metadata were prepared.

### Liver protein isolation and processing for proteomics experiment

Frozen liver tissue (100 mg) was sliced into thin pieces and transferred to a bead-beating tube containing 5 zirconia beads (2 mm) and 500 µl lysis buffer (20 mM HEPES, 100 mM NaCl, 0.05% Triton X-100, 1 mM DTT). Bead beating was carried out in an MP Biomedical Bead beating homogenizer with 20 sec on/1 min off twice. The tissue lysate was clarified by centrifuging at 12,000 g, 15 min at 4°C, and the supernatant was recentrifuged at 12,000 g, 10 min at 4°C. The amount of protein in the tissue lysates was quantified by using the Bicinchoninic Acid assay (BCA) method.

An equal amount of liver protein (100 µg) from each biological replicate was dried in a SpeedVac (Labconco). After resuspending it in TEAB (100 mM, 100 µl), the proteins were reduced by incubating with TCEP (200 mM, 5 µl) for 1 hour at 55°C. Then, these proteins were alkylated by incubating with iodoacetamide (375 mM, 5 µl) at room temperature for 30 min in the dark. These proteins were precipitated using pre-chilled acetone (6:1, v/v) and by incubating overnight at -20°C. These mixtures were centrifuged at 8,000 g for 10 min at 4°C. The pellet was resuspended in TEAB (100 mM, 100 µl) and incubated overnight with sequencing-grade trypsin (2.5 µg) at 37°C. The digestion was stopped by the addition of 41 µl of anhydrous acetonitrile, and the tryptic peptides were dried in a SpeedVac (Labconco) at 40°C. The dried peptides were resuspended in 300 µl of 0.1% trifluoroacetic acid and desalted using Pierce™ Peptide Desalting Spin Columns by following the manufacturer’s instructions. The eluted peptides were dried in a SpeedVac at 40°C and stored for LC-MS/MS data acquisition.

### Liver proteomics data acquisition using LC-MS/MS and analysis

The dried tryptic peptides were resuspended in 0.1% Formic Acid (Merck, Cat no. 5330020050) in MS-grade water (Fisher Scientific, Cat no. AAB-W6-4). One µg of tryptic peptides were loaded and separated first on PepMap Neo C18 5 µm 300 µm x 5 mm 1500 bar Trap column (Thermo Scientific, Cat no. 174500) and then on EASY-Spray PepMap Neo C18 Column, 2 µm, 75 µm x 500 mm, 1500 bar (Thermo Scientific, Cat no. ES75500PN) using Thermo Scientific Vanquish Neo UHPLC system at a flow rate of 300 nl/minute using a gradient of solvent A (100% H_2_0+0.1% FA) and solvent B (80% ACN+0.1% FA). The peptides were separated using a reversed-phase liquid chromatography using a gradient from 5% to 50% of solvent B over 120 minutes, with a flow rate of 300 nl/min. Further, the MS1 data were acquired in positive ion mode using an Orbitrap with a resolution of 60,000 and a mass range from 350 to 1200 m/z. Precursor ions were fragmented using higher-energy C-trap dissociation (HCD) in an ion trap (IT) with a collision energy of 30% in a data-dependent mode, accompined by a rapid IT scan rate. Precursor ions with +2 to +7 charge and monoisotopic ions were selected. Parent ions, once fragmented, were excluded for 60s with an exclusion mass width of ± 10 ppm.

The raw MS/MS data were searched against the UniProt proteome database of *Mus musculus* (UP000000589; 54690 sequences on 2025_06_06) at an FDR of 5% using Proteome Discoverer software version 3.1.0.638 (Thermo Scientific). A maximum of two trypsin cleavages was allowed with a precursor mass tolerance of 10 ppm and fragment mass tolerance of 0.6 Da. Carbamidomethylation (+57.021 Da) of cysteine residues at the C-terminal was selected as the static modification, and oxidation of methionine residues (+15.995 Da) and acetylation at the N-terminus (+42.011 Da) were selected as the dynamic modifications. Peptides and master proteins identified with high confidence were annotated, and their quantitative abundances were normalized to the same total peptide amount. The final matrix was used for statistical analysis, and proteins with a log_2_ fold change of ≥ ±1.0 and a -log_10_ adjusted p-value of ≥ 1.3 were classified as deregulated proteins.

### Mtb H37Rv infection

A set of mice completing 30 days of TRF and a control ALF group were aerosol infected with a low dose (100-120 colony-forming units; CFU) of animal-passaged Mtb H37Rv using a Madison chamber in the tuberculosis aerosol challenge facility (BSL-III) in ICGEB. Mice were humanely euthanized on days 1 and 21 post Mtb H37Rv infection to monitor their tissue mycobacterial burden. During this period, mice infected with Mtb from both the TRF and ALF groups had access to food for 24 hours. Mice tissues (lung, spleen, and liver) were aseptically collected and homogenized in phosphate-buffered saline, and the tissue homogenates were plated on 7H11 agar plates supplemented with PANTA. The plates were incubated at 37°C in a humidified incubator for 21 days, and the colonies were enumerated to calculate the colony-forming units (CFUs).

### Tissue immune cell phenotyping

Harvested tissues (lungs, spleen) of experimental mouse groups were incubated with collagenase D (1 µg/mL) and DNase I (0.5 mg/mL) in RPMI 1640 medium at 37°C for 30 min. The cell suspension was passed through a cell strainer (70 µm) and centrifuged at 400 g at 4°C for 5 min. The cell pellet was lysed with RBC cell lysis buffer at room temperature for 5 min. After washing the immune cells with PBS, they were stained with viability dye (LIVE/DEAD™ Fixable Near-IR Dead Cell Stain) and surface markers (CD3-FITC and CD4-BV785).

For immune cell isolation from bone marrow, the epiphyses of the femur and tibia were cut with scissors and placed in a 0.6 mL microcentrifuge tube (MCT), then kept in a separate tube. After centrifuging at 2,000 rpm for 5 sec, the cell pellet was collected, and then resuspended in RPMI 1640, the medium was centrifuged again. The pellet was resuspended in RBC cell lysis buffer after incubating for 5 min at room temperature, and the cells were washed with PBS before staining with viability dye (LIVE/DEAD™ Fixable Near-IR Dead Cell Stain) and surface markers (CD3-FITC and CD4-BV785) on ice for 30 min. After washing, the cells were washed with PBS and resuspended in FACS buffer (0.5% FBS in PBS) before data acquisition using BD LSRFortessa X-20, and data were analyzed using FCS Express Version 6.0.

### Cytokine estimation

Serum cytokine levels were measured using the LEGENDplex Mix and Match mouse inflammation panel, following the manufacturer’s instructions (BioLegend). The data were acquired using BD LSRFortessa X-20, and the data were analysed using web-based LEGENDplex™ Data Analysis Software Suite.

### Data analysis

All group-specific data were presented as mean ± standard error of the mean (sem). Mann-Whitney or Welch’s t-test was used to compare statistical significance in parameters between the two groups, and a p-value<0.05 at a 95% confidence interval was considered statistically significant. The iPGTT data were analyzed using two-way analysis of variance (ANOVA). The final data matrix generated from metabolomics and proteomics data was uploaded to MetaboAnalyst (6.0), and the data were log-transformed and auto-scaled for multivariate data analysis. Metabolites or proteins with a log_2_ fold change of ≥ ±1.0 and a log10-adjusted p-value of ≥ 1.3 were classified as deregulated molecules. The plots were generated using GraphPad Prism Version 8 or the R package “ggplot2”.

## Results

### TRF influences body weight gain and impacts host glucose homeostasis

To monitor the effect of TRF on the host, 6–8 weeks C57BL/6 mice were subjected to a TRF regimen with access to food for 8 h from ZT13 to ZT21, whereas the control *ad libitum* (ALF) group had unrestricted access to food (Figure 1A). The TRF group consumed a similar amount of food to the ALF group (Figure 1B) and, in the initial two weeks, lost 1.5% of body weight (Figure 1C). After 30 days of TRF, the mice lost 5.49% of body weight, whereas the ALF control group gained 16.62% of body weight (Figure 1C). At days 7 and 14, mice undergoing TRF lost significant body weight compared to the ad-libitum controls (Supplementary Figure S1A).

**Figure 1.**
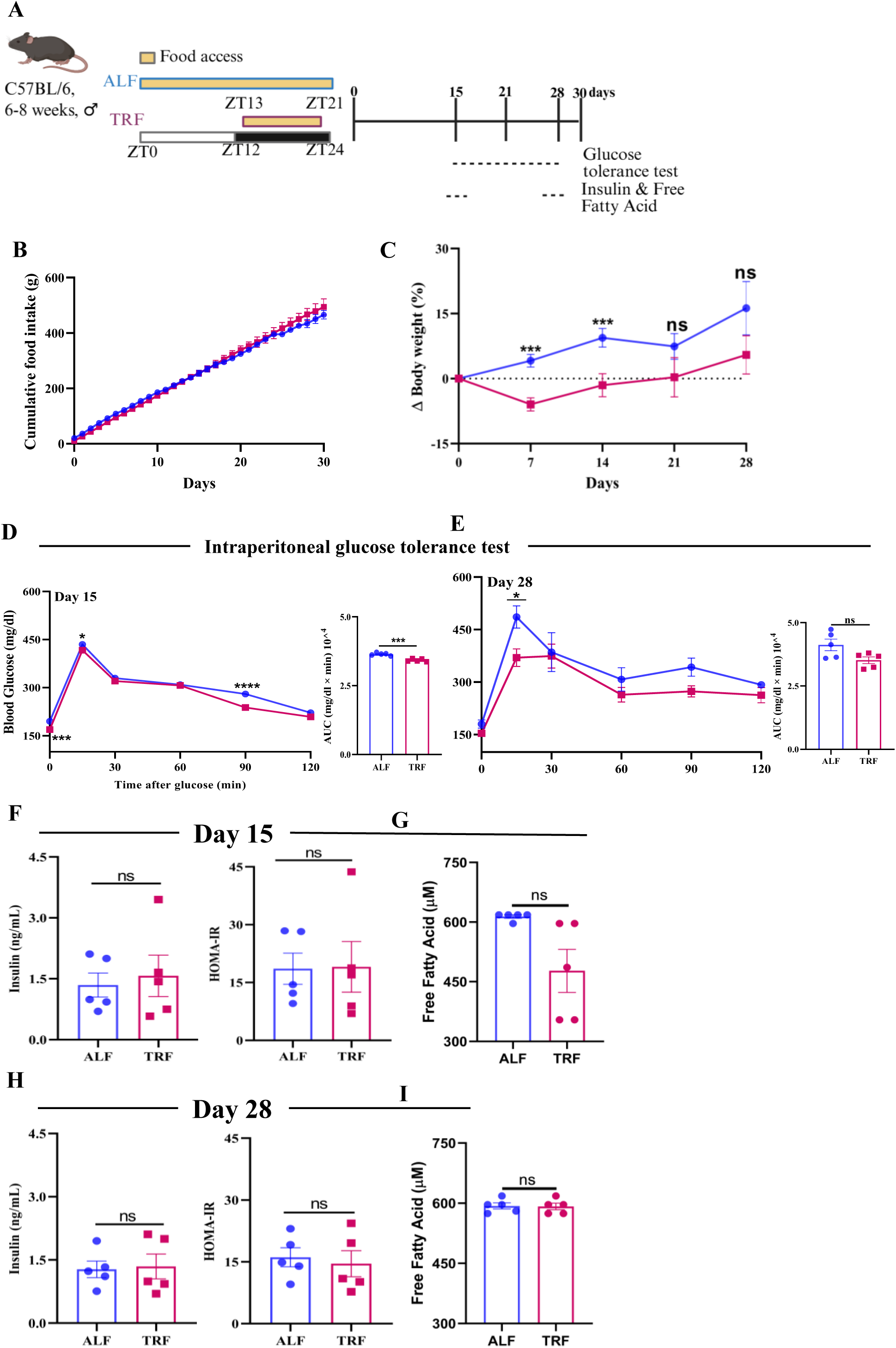
TRF alters the glucose homeostasis at early time points but does not affect fasting serum insulin and free fatty acid levels. **A**. Schematic of the experimental design adopted in the study. **B**. Cumulative food intake per cage in grams. **C**. Change in body weight with time. **D**. and **E**. Intraperitoneal glucose tolerance test at Days 15 and 28 post-TRF. The quantification of the AUC above baseline is shown in the inserts. **F**. and **H**. Insulin levels after 16 hr of fasting and corresponding HOMA-IR levels at Day 15 and Day 28, respectively. **G**. and **I**. Free fatty acid levels after 16 hr of fasting at Day 15 and Day 28, respectively. The number of mice analyzed per group for each figure was 5, except for **B,** for which 2 cages per group were analyzed. Each dot represents a biological replicate. Statistical significance was determined using Holm-Sidak method with correction for multiple comparison in **B**. and **C**., two-way repeated-measures ANOVA and Sidak’s multiple comparisons tests for ipGTT and unpaired t-test with Welch’s correction for AUC in **D.** and **E**. and unpaired t-test with Welch’s correction in **F., G., H. and I.** *p≤0.05, ***p≤0.001, ****p≤0.0001, ns = not significant.

On day 14, the TRF group had a significantly lower area under the curve for the intraperitoneal glucose tolerance test (Figure 1D). At days 21 and 28, both the TRF and ALF groups had similar areas under the curve for the intraperitoneal glucose tolerance test, indicating similar glucose tolerance (Figure 1E and Supplementary Figure S1B). Fasting serum insulin and free fatty acid levels and HOMA-IR between the TRF and ALF mice were similar at days 15 and 28 (Figure 1F to I). Initiation of TRF resulted in significant weight loss at initial time points (up to day 14), but at later time points, body weight was similar between the ALF and TRF groups. Similar observations, like minimal body weight gain and changes in glucose homeostasis, have been reported in humans undergoing TRE.^1^

### TRF alters amino acid and fatty acid metabolism

We performed a global metabolomics experiment to monitor TRF-induced metabolic alterations in the circulation and liver, if any (Figure 2A). We selected serum to monitor systemic changes, and the liver, given its critical contribution as a metabolic hub. Principal component analysis (PCA) of the serum metabolome showed minor overlap between the TRF and ALF groups (Figure 2B). A set of 1,123 metabolites was identified in the serum, out of which 51 were deregulated (log_2_ fold change ≥ ±1.0; -log_10_ p-value >1.3) (Supplementary Table S1). These deregulated molecules originated from the metabolism of amino acids (alanine, aspartate, glutamate, arginine, proline, tyrosine), lipid biosynthesis including the elongation and degradation of fatty acids), pyrimidine metabolism, and steroid hormone biosynthesis pathways (Figure 2C and D). Serum palmitoleic acid levels were significantly low, and higher cytidine and cystine levels were observed in the TRF groups, indicating an altered fatty acid and pyrimidine metabolism between the study groups (Supplementary Figure S2A). Similarly, global liver metabolome data showed slight overlap in the PCA plot (Supplementary Figure S2B and S2C). Through GC-MS-based metabolite analysis, we captured 119 liver metabolites, out of which eight (propanoic acid, methylmaleic acid, gluconic acid, lysine, benzaldehyde, 10-undecynoic acid, arabinonic acid, and ribonolactone) were dysregulated (log_2_ fold change ≥ ±1.0; p-value <0.05) in the TRF mice (Supplementary Table S2). These molecules are involved in biotin metabolism, propanoate metabolism, the pentose phosphate pathway, and lysine degradation pathways, indicating significant differences in the liver metabolic phenotypes of the TRF and ALF groups (Supplementary Figure S2D and S2E).

**Figure 2.**
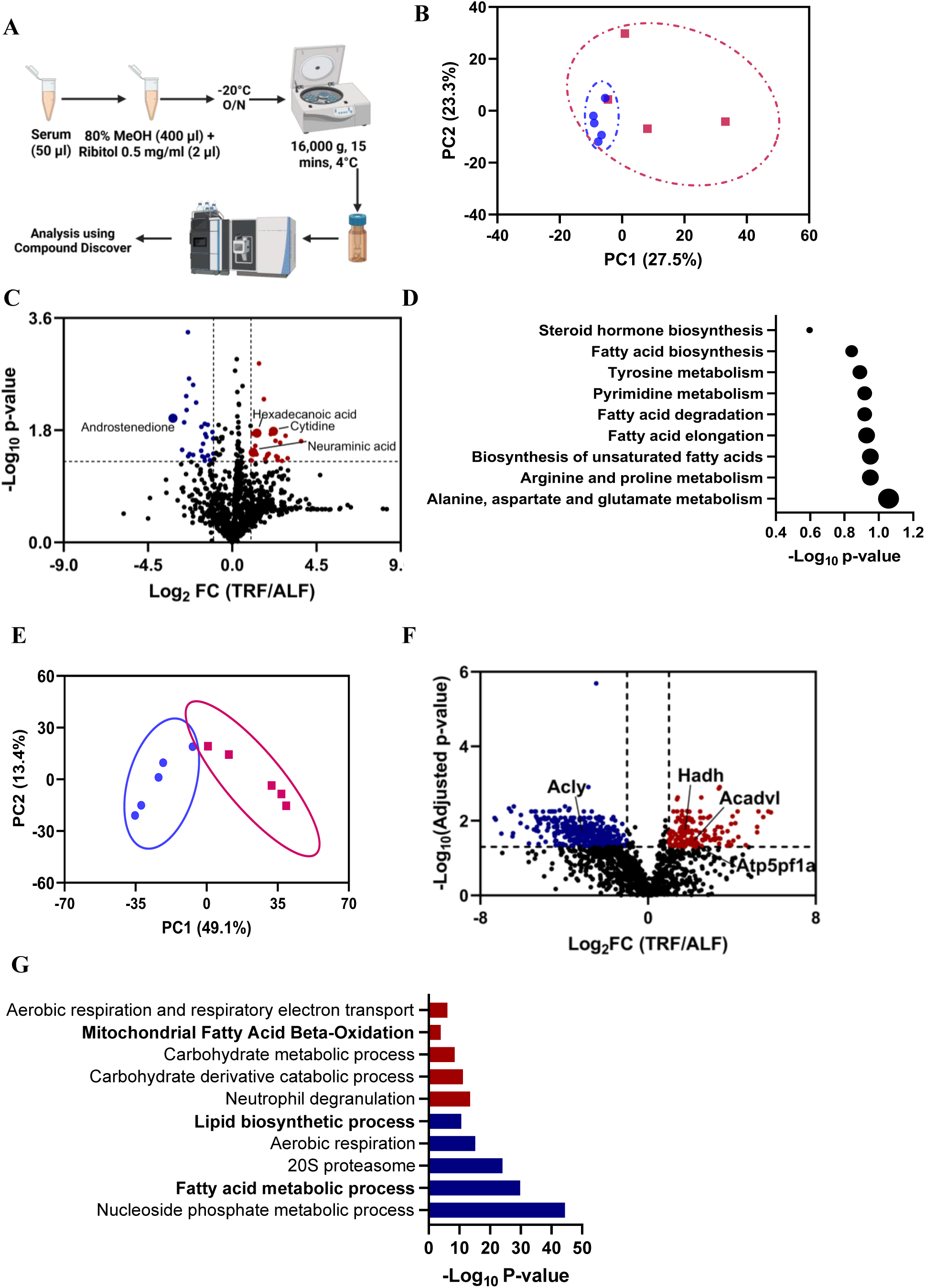
TRF-induced metabolism remodelling was observed at both tissue and systemic circulation levels. **A.** Schematic representation of the method used for metabolite profiling from the TRF and ALF mice serum using LC-MS. **B.** Principal component analysis (PCA) of the serum metabolites between the TRF and ALF mice. **C.** Volcano plot showing the deregulated metabolites in the serum of the TRF and ALF mice. **D.** Metabolite set enrichment analysis (MSEA) of the deregulated metabolites. The size of the dot represents the enrichment ratio. **E.** PCA of the liver proteomics between the TRF and ALF mice. **F.** Volcano plot showing the deregulated proteins in the liver of the TRF and ALF mice. **G.** Pathway enrichment analysis of the deregulated proteins. Upregulated pathways are shown in red, whereas the deregulated pathways are shown in blue.

TRF also majorly impacted the liver proteome and showed a clear separation between the ALF and TRF mice (Figure 2E). TRF mice liver showed decreased abundance of 437 proteins (log_2_ fold change ≥ -1.0; p-value <0.05), and a set of 152 proteins had higher abundance (log_2_ fold change ≥ 1.0; p-value <0.05; Figure 2F, Supplementary Table S3). The pathway enrichment analysis using the deregulated liver proteome data showed upregulated fatty acid beta-oxidation and downregulated lipid biosynthetic process (Figure 2G). Apart from the increased utilization of fatty acids, TRF mice had higher abundance of proteins such as ATP synthase F(1) complex subunit alpha, Cytochrome c oxidase subunit 5A and Glyceraldehyde-3-phosphate dehydrogenase involved in aerobic respiration and electron transport chain. We performed global correlation analysis on the significantly deregulated liver metabolites and the top 20 down/up-regulated proteins. We observed a significant positive correlation between methylmaleic acid, propanoic acid and lysine, with Hao1 involved in fatty acid alpha-oxidation (Supplementary Figure S2F). Thus, 30 days of continuous 8-hour TRF in mice resulted in significant alterations in amino and fatty acid metabolism at both systemic and tissue levels.

### Pre-exposure to TRF for 30 days minimally affected tissue mycobacterial burden at 21 days post-infection (dpi)

To determine if 30 days of TRF improved the host’s ability to handle respiratory infections such as Mtb, C57BL/6 mice completing 30 days of TRF were aerosol infected with 100–400 CFU of animal passaged Mtb H37Rv (Figure 3A and Supplementary Figure S3A). After MTB infection, the TRF mice group received food *ad libitum,* similar to the ALF group. The Mtb-infected TRF and ALF control mice showed lower body weights compared to their pre-infection body weights, and body weight was similar between the groups at 21 dpi (Figure 3B and Supplementary Figure S3B). At 21 dpi, random blood glucose levels between the Mtb-infected TRF and ALF mice were similar (Supplementary Figure S3C). The gross tissue pathology of the TRF-TB and ALF-TB mouse groups showed similar pathology (Figure 3C). At 21 dpi, the Mtb-infected TRF and ALF mice groups had similar lung and spleen tissue mycobacterial burden (Figure 3D). Therefore, pre-exposure to TRF for 30 days did not benefit the C57BL/6 mice in terms of tissue mycobacterial clearance at the early time points, i.e., 21 dpi. However, TRF mice infected with Mtb may have impacted the tissue immune system.

**Figure 3.**
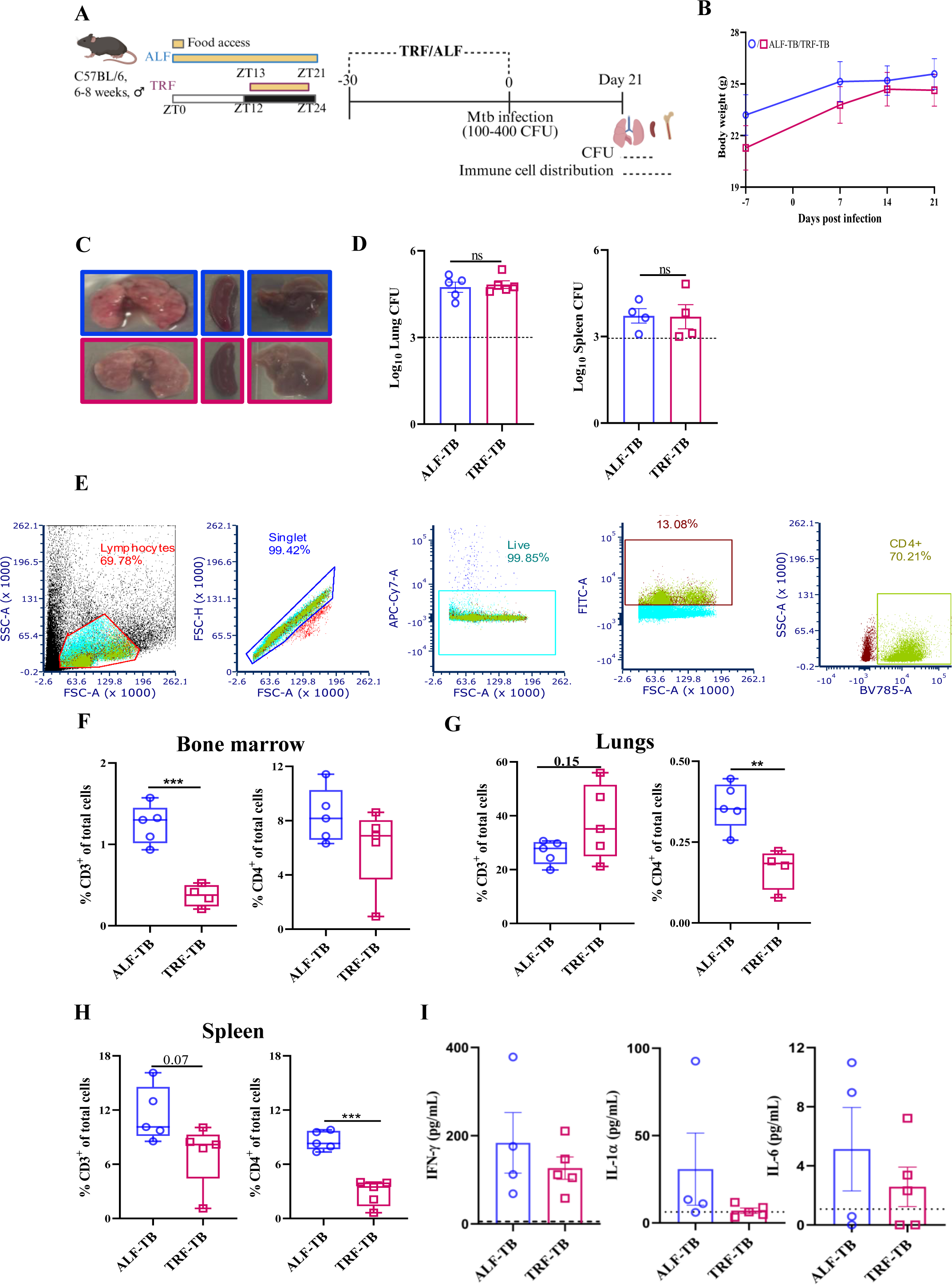
TRF decreases the CD4^+^ T cell number in a tissue-dependent manner at 21 days post-Mtb H37Rv infection (dpi). **A.** Schematic of experimental design adopted in the study. **B.** Change in body weight with time after *Mycobacterium tuberculosis* (Mtb) infection. **C.** Gross pathology of the lungs, spleen and liver of the ALF and TRF mice. **D.** Mycobacterial burden in the lungs and spleen at 21 dpi. **E.** The gating strategy was adopted to determine CD3^+^ T cells and CD3^+^ CD4^+^ T cells. Percentage of CD3^+^ and CD3^+^ CD4^+^ T cells in (**F.**) bone marrow, (**G.**) lungs and (**H.**) spleen. **I.** Levels of pro-inflammatory cytokines IL-1α, IFN-γ and IL-6 in the serum. Each dot represents a biological replicate. Statistical significance was determined using an unpaired t-test with Welch’s correction. **p≤0.01 and ***p≤0.001.

### Mtb-infected mice pre-exposed to TRF showed differential tissue immune cell distribution

To monitor the tissue immune cell distribution of these Mtb-infected TRF and control ALF mice groups, we monitored the distribution of CD3^+^ and CD4^+^ T cells in multiple tissues (bone marrow, lungs, and spleen; Figure 3E) at 21 dpi. The bone marrow of TRF-TB mice had significantly lower CD3^+^ and CD4^+^ T cells compared to the ALF-TB group (Figure 3F and Supplementary Figure S4A). Similarly, the lung CD4^+^ T cell population was significantly lower in the TRF-TB group (Figure 3G and Supplementary Figure S4B). The frequencies of splenic CD3^+^ and CD4^+^ T cells were similar between the study groups (Figure 3H and Supplementary Figure S4C). The circulatory IL-6, IFN-γ, and IL-α levels between Mtb-infected TRF and ALF mice groups at 21 dpi were similar (Figure 3I). Thus, pre-exposure to TRF resulted in decreased numbers of CD3^+^ and CD4^+^ T cells, key immune cell types for host defence against Mtb, with similar mycobacterial load at the early time point, i.e., 21 dpi.

### TRF-induced metabolic changes persisted in TRF-TB mice

To determine whether TRF-induced metabolic alterations in circulation and liver persist after TRF is discontinued post-Mtb infection, an untargeted metabolomics experiment was conducted (Figure 4A). The PCA plot of the serum metabolome data showed a clear separation between the TRF-TB and ALF-TB groups (Figure 4B). Out of the identified 1120 serum metabolites, 22 were deregulated (log_2_ fold change ≥ ±1.0; p-value < 0.05) in the TRF-TB group (Supplementary Table S4). Pathway enrichment analysis of the deregulated metabolites identified in serum, revealed perturbed nicotinate and nicotinamide metabolism, sphingolipid metabolism, and fatty acid and steroid hormone biosynthesis (Figure 4C and D). PCA plot of the liver metabolome data of TRF-TB and ALF-TB showed a slight overlap (Supplementary Figure S5A). Out of the 121 identified metabolites in the liver, the TRF-TB mice group showed 14 deregulated (log_2_ fold change ≥ ±1.0; p-value <0.05) metabolites (Supplementary Figure S5B and Supplementary Table S5). Pathway enrichment analysis of deregulated metabolites showed perturbed linoleic acid metabolism, fructose and mannose metabolism, galactose metabolism, biosynthesis of unsaturated fatty acids, tyrosine metabolism, amino acid and nucleotide sugar metabolism, and arachidonic metabolism (Supplementary Figure S5B and S5C). Therefore, it is essential to note that TRF-induced metabolic changes persisted in the TRF mice following Mtb infection.

**Figure 4.**
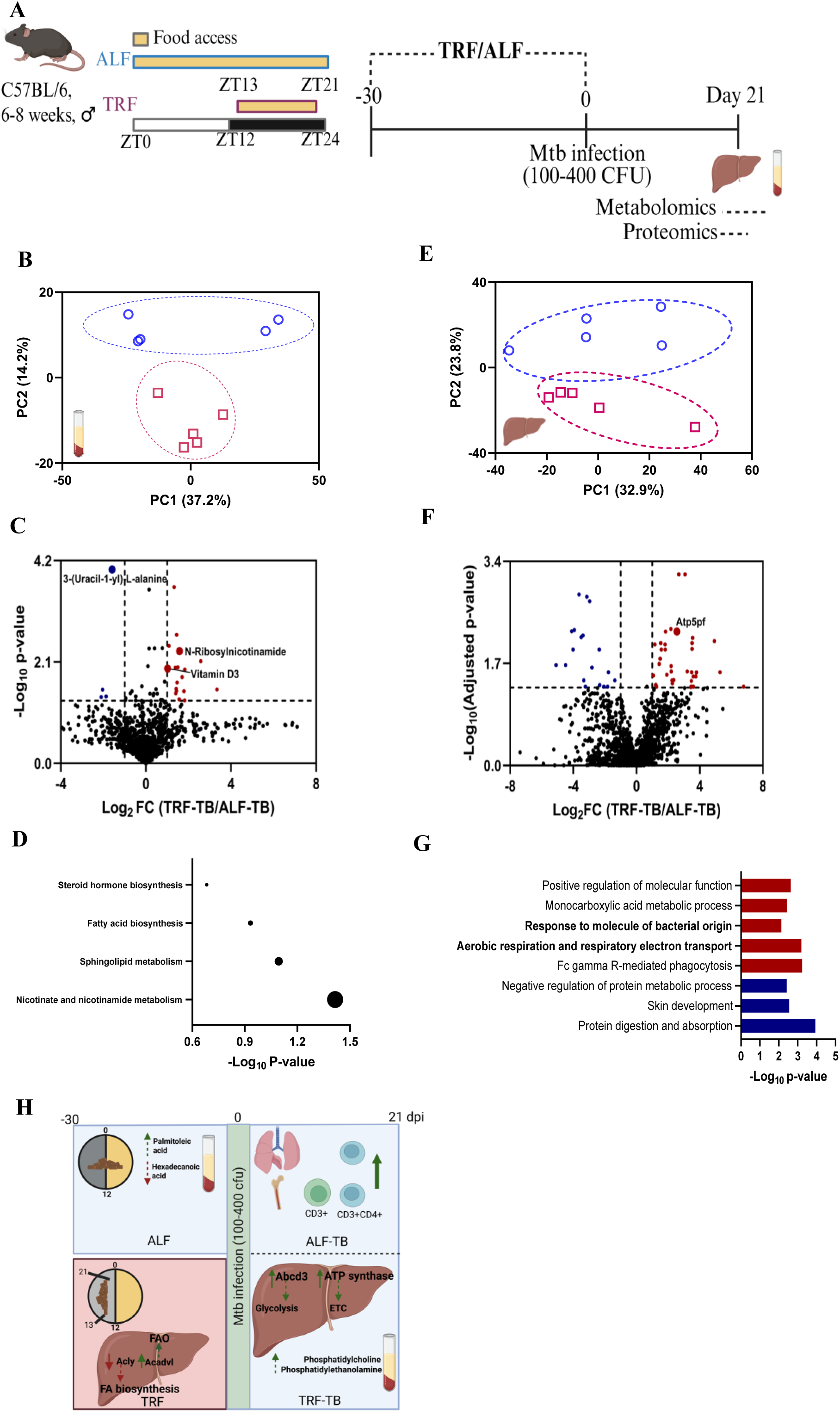
TRF-induced metabolic changes persist post 21 days of Mtb infection. **A.** Schematic of experimental design adopted in the study. **B.** Principal component analysis (PCA) of the serum metabolites between the TRF-TB and ALF-TB mice. **C.** Volcano plot showing the deregulated metabolites in the serum of the TRF-TB and ALF-TB mice. **D.** Metabolite set enrichment analysis (MSEA) of the deregulated metabolites. The size of the dot represents the enrichment ratio. **E.** PCA of liver proteomic analysis between the TRF-TB and ALF-TB mice. **F.** Volcano plot showing the deregulated proteins in the liver of the TRF-TB and ALF-TB mice. **G.** Pathway enrichment analysis of deregulated proteins. Upregulated pathways are shown in red, whereas the deregulated pathways are shown in blue. **H.** Effect of TRF on the host metabolism and the immune system.

Similar to metabolic analysis, we investigated whether TRF-induced changes in the liver proteome persist after Mtb infection. The PCA plot highlighted the difference in the liver proteome of the ALF-TB and TRF-TB mice (Figure 4E). TRF-TB mice had a set of 56 deregulated liver proteins (log_2_ fold change ≥ ±1.0; adjusted p-value < 0.05; Figure 4F, Supplementary Table S6). The pathway enrichment analysis showed significant upregulation of metabolic pathways (aerobic respiration and respiratory electron chain, and monocarboxylic acid metabolic process) and immune system-related pathways (response to molecules of bacterial origin and Fc gamma R-mediated phagocytosis; Figure 4G). We performed global correlation analysis on the significantly deregulated liver metabolites and the top 20 down/up-regulated proteins of the ALF-TB vs TRF-TB mice. We observed a significant positive correlation between proteins involved in the electron transport chain (Atp5f, Cox5a, Sdhaf4 (Supplementary Figure S5D). Thus, alterations induced by TRF in amino acid and fatty acid metabolism persist even after its discontinuation.

## Discussion

In this study, we investigated the effect of TRF on the host immune system, specifically in the context of infection with the intracellular pathogen *Mycobacterium tuberculosis*. TRF (or TRE) is a dietary intervention that reduces the food consumption window (a minimum of less than 12 hours) without altering the quality or quantity of food consumed. This has been shown to have pleiotropic metabolic benefits, especially in the context of lifestyle-induced metabolic diseases.

TRF in mice consuming a high-fat diet (HFD) demonstrated the ability to prevent weight gain, improve glucose homeostasis, and reduce the size of lipid droplets in adipose tissue.^2^ These effects are achieved despite taking similar calories and primarily by altering the diurnal expression of metabolic genes, i.e., matching food availability with the peak of expression of metabolic enzymes.^17^ Although TRF has been demonstrated to have pleiotropic health benefits, translation of these benefits to humans has shown variable efficacy.^12,13^

We observed that a 30-day treatment with TRF in C57BL/6 male mice resulted in significant body weight loss during the initial period, which was not observed at later time points and subsequently became similar. That corroborates with earlier reports that the effect of TRF on lowering body weight was demonstrated only at early time points. Multiple reports have shown mixed findings on the loss of body weight in humans from various ethnicities who adopted TRE for specified time periods. Most reports showed minimal body weight loss in humans undergoing TRE.^1,14–16^ Mice receiving HFD following TRF show significantly lower body weight gain compared to ALF controls.^2,17^ In our study, mice had access to a normal chow diet in which only 13% of calories are derived from fat, whereas in HFD, fat is responsible for 60% of calories. We observed better glucose homeostasis in TRF mice at an early time (day 15 of TRF), but the effect was abrogated by day 28. During this period, we did not observe any differences in fasting insulin levels or HOMA-IR between the TRF and ALF mice. This could be due to a lack of insulin resistance in 6–8 weeks old male C57BL/6 mice being fed a normal chow diet, in contrast to those fed with HFD. TRF has been reported to reduce insulin resistance and improve HOMA-IR levels in diabetic individuals and mice consuming HFD.^1,2^ The serum fasting free fatty acid levels in the TRF mice at days 15 and 28 were similar.

TRE in humans is reported to influence pathways associated with amino acids and lipid metabolism.^16^ We also observed that TRF remodeled the host metabolome in the circulation and metabolically active tissue, such as the liver. Pathways related to amino acid metabolism (alanine, aspartate and glutamate metabolism, arginine and proline metabolism, and tyrosine metabolism), fatty acid metabolism (biosynthesis of unsaturated fatty acids, fatty acid elongation, fatty acid degradation, fatty acid biosynthesis, and steroid hormone biosynthesis), and pyrimidine metabolism were enriched in the circulation of mice completing 30 days of TRF. Pathways associated with biotin metabolism, propanoate metabolism, pentose phosphate pathway and lysine degradation were enriched in the liver of TRF mice. The liver proteome of TRF mice indicated a perturbed fatty acid metabolism. The pathway enrichment analysis indicated increased reliance on oxidative phosphorylation and fatty acid metabolism for host energetic requirements. Fatty acid and amino acid metabolism-related pathways remain altered at 21 dpi even though TRF is discontinued prior to Mtb infection (Figure 4H). Very few studies have investigated the effect of TRF on the host immune system in the context of bacterial infection. However, caloric restriction (CR) has been shown to significantly alter the number and functionality of immune cells. CR is reported to significantly reduce the spleen mycobacterial burden at 40 dpi with no effect on the lung mycobacterial burden in C57BL/6 mice intravenously infected with Mtb H37Rv.^19^ CR also resulted in a significant decrease in the IFN-γ levels in the lung homogenate.^19^ In this study, we observed similar lung or spleen mycobacterial burden in the TRF mice at 21 dpi. We assessed both the tissue mycobacterial burden and immune cell distribution at 21 dpi, as the adaptive immune response peaks and the Mtb levels reach a plateau in C57BL/6 mice by 21 dpi.^20^ The circulatory pro-inflammatory cytokines (IL-1α, IFN-γ, and IL-6) levels between the Mtb-infected TRF and control study groups (TRF-TB and ALF-TB) were similar. At 21 dpi, the TRF mice’s bone marrow had a lower CD3^+^ and CD4^+^ T cell population and lower levels of CD4^+^ T cells in the lungs. CD4^+^ T cells play a crucial role in the host immune response against Mtb, but a decrease in the number of CD4^+^ T cells did not compromise the containment of Mtb infection by the TRF mice. Earlier reports stated that CR initiated before the Mtb infection resulted in a decreased number of CD3^+^CD4^+^ T cells in the lungs but not in the spleen of intravenously Mtb H37Rv-infected DBA/2 mice at 40 dpi.^19^ Differential effect of TRF and CR on tissue mycobacterial burden and immune cell distribution could be explained by differences in the energetics, in the mouse strain (C57BL/6 vs DBA/2), differences in timepoints (21 dpi vs 40 dpi) selected for sacrifice, and differences in the mode of infection (aerosol vs intravenous).^19^

A limitation of our study is that we did not assess how TRF affects the functionality of CD3^+^ or CD4^+^ T cells or the distribution of other immune cells (such as macrophages and neutrophils) involved in the host defence against Mtb. Also, the current study did not have a group in which TRF was continued throughout Mtb infection or initiated after Mtb infection. The inclusion of these groups and maintaining them for a longer period could help determine whether continued TRF or TRF initiated after Mtb infection might have a positive effect on the immune system.

In conclusion, our study demonstrated that TRF for 30 days remodelled host metabolic phenotype by altering amino acid and lipid metabolism, and these beneficial effects persist even if TRF is discontinued. However, discontinuing the TRF had no additional effect on mycobacterial clearance at 21 dpi. Pre-exposure to TRF affected both tissue-level metabolism and the distribution of immune cells (CD3^+^ and CD4^+^ T cells) in Mtb-infected mice. Since the decrease in the number of CD3^+^ and CD4^+^ T cells did not compromise the host’s mycobacterial clearance ability, it indicates that pre-exposure to TRF might improve the functionality of CD3^+^ and CD4^+^ T cells.

## Supporting information

This article contains supporting information.

## Acknowledgements

We acknowledge the Department of Biotechnology (DBT), Government of India, for supporting activities through research grants and supporting the Tuberculosis Aerosol Challenge Facility at ICGEB, New Delhi and the ICGEB, New Delhi, for providing core support to RKN. AG and NY received the Junior Research Fellowship from DBT, Government of India. The authors are thankful to Ms Suchitra Jena for her help in animal experiments.

