## Supplemental Information for "Pre-exposure to time-restricted feeding reduces tissue CD4^+^ T cells with a limited effect on *Mycobacterium tuberculosis* clearance at the early time point"

### Slide 1
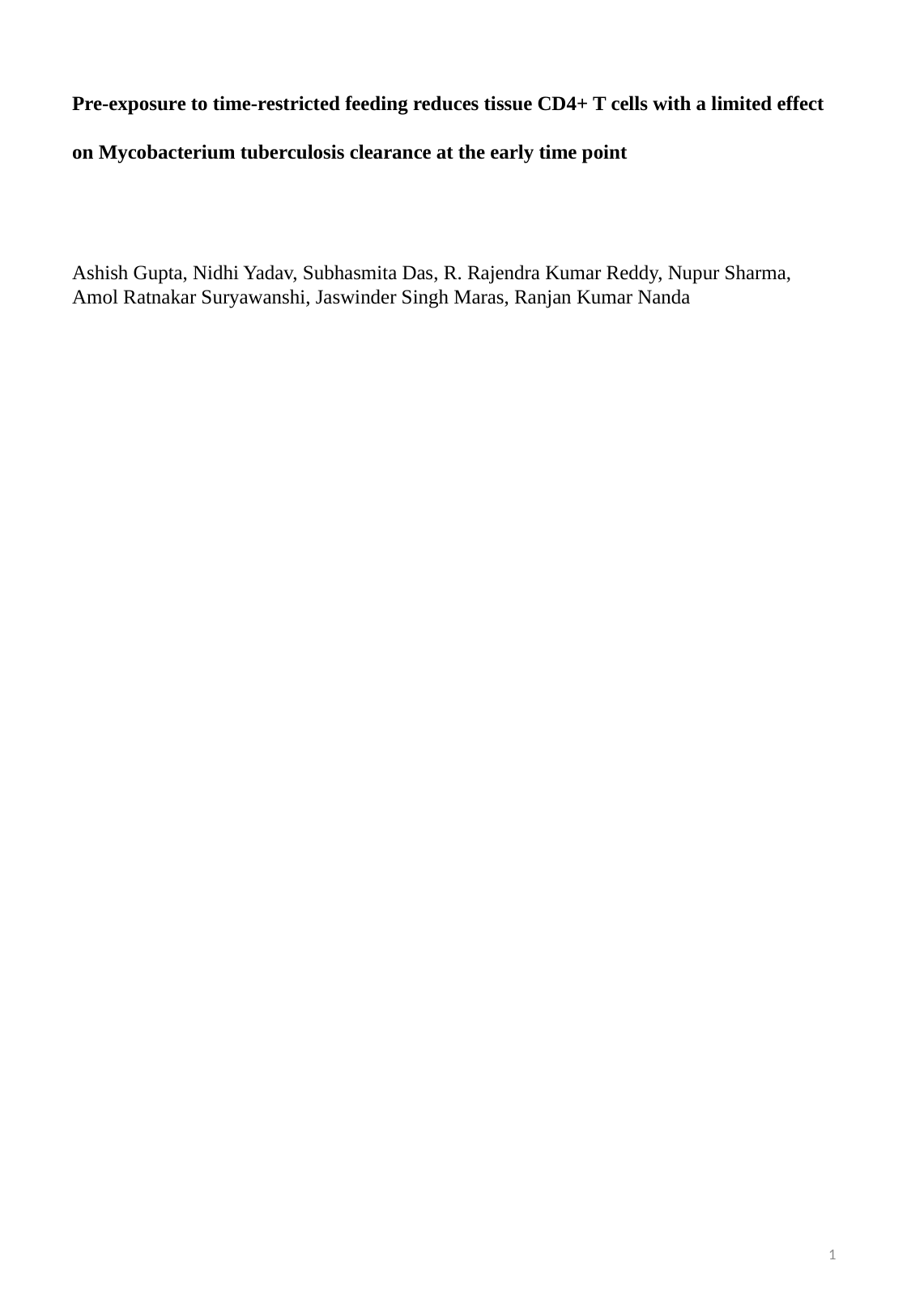

Pre-exposure to time-restricted feeding reduces tissue CD4+ T cells with a limited effect on Mycobacterium tuberculosis clearance at the early time point
Ashish Gupta, Nidhi Yadav, Subhasmita Das, R. Rajendra Kumar Reddy, Nupur Sharma, Amol Ratnakar Suryawanshi, Jaswinder Singh Maras, Ranjan Kumar Nanda
1

### Slide 2
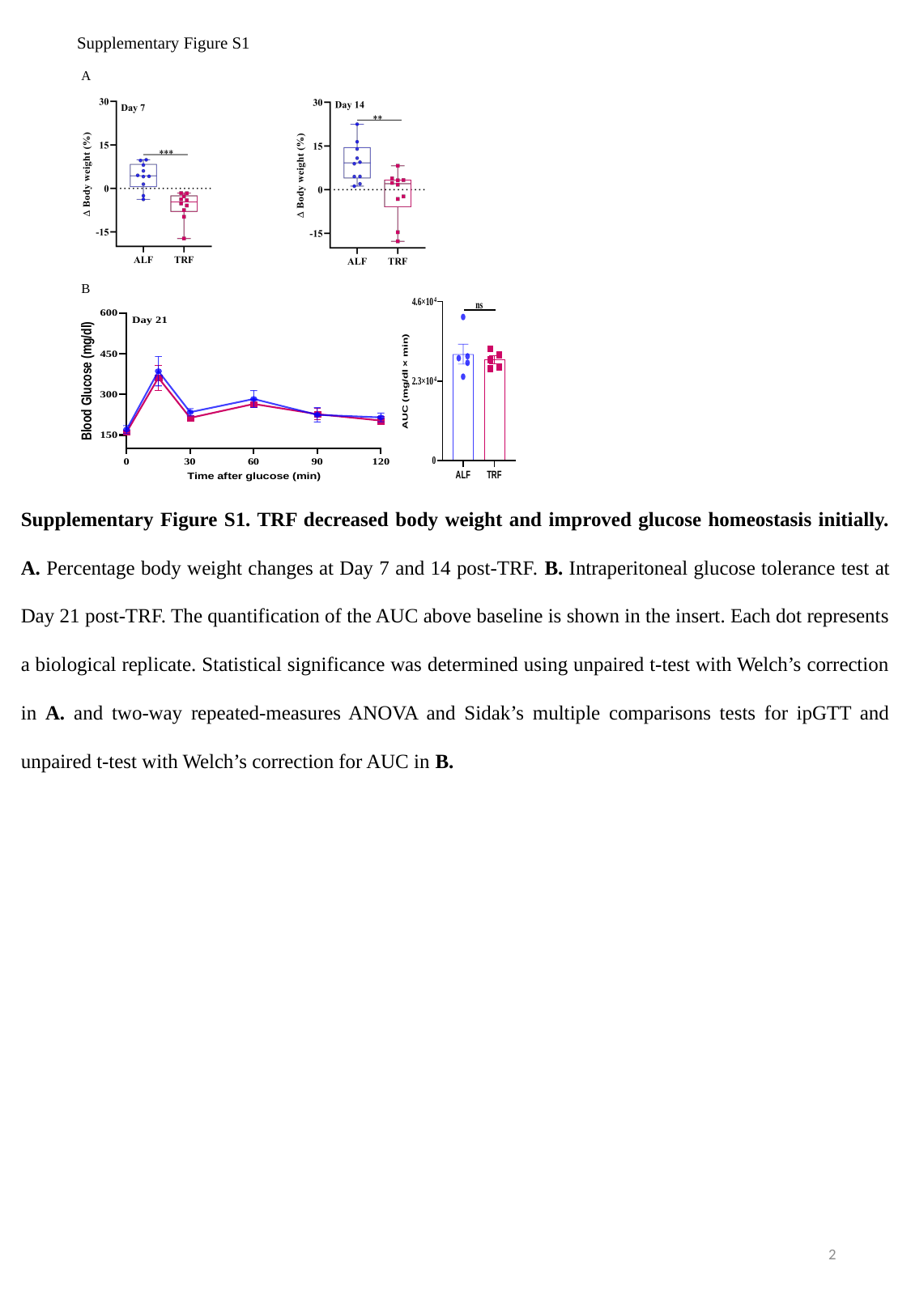

Supplementary Figure S1
A
B
Supplementary Figure S1. TRF decreased body weight and improved glucose homeostasis initially. A. Percentage body weight changes at Day 7 and 14 post-TRF. B. Intraperitoneal glucose tolerance test at Day 21 post-TRF. The quantification of the AUC above baseline is shown in the insert. Each dot represents a biological replicate. Statistical significance was determined using unpaired t-test with Welch’s correction in A. and two-way repeated-measures ANOVA and Sidak’s multiple comparisons tests for ipGTT and unpaired t-test with Welch’s correction for AUC in B.
2

### Slide 3
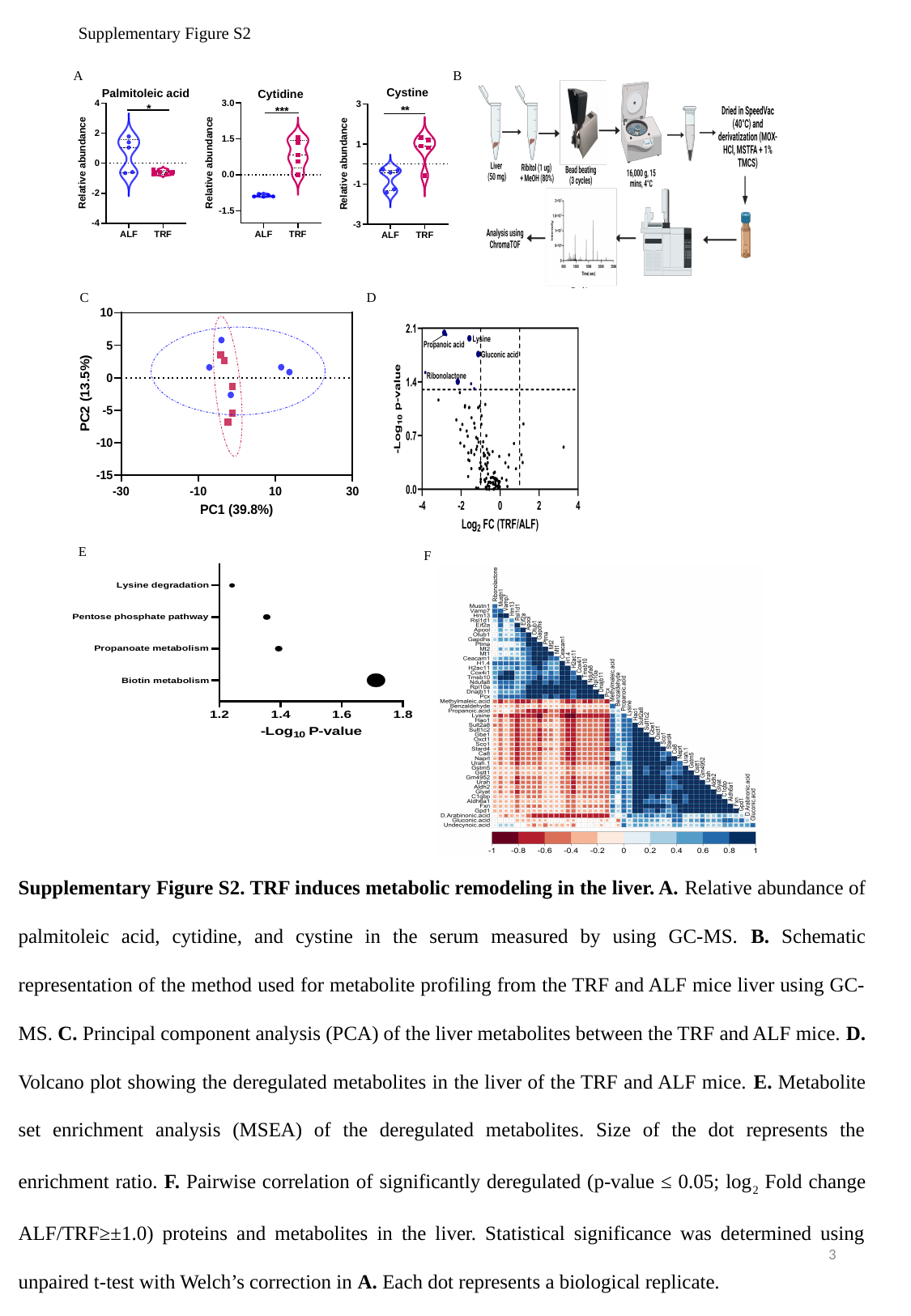

Supplementary Figure S2
A
B
C
D
E
F
Supplementary Figure S2. TRF induces metabolic remodeling in the liver. A. Relative abundance of palmitoleic acid, cytidine, and cystine in the serum measured by using GC-MS. B. Schematic representation of the method used for metabolite profiling from the TRF and ALF mice liver using GC-MS. C. Principal component analysis (PCA) of the liver metabolites between the TRF and ALF mice. D. Volcano plot showing the deregulated metabolites in the liver of the TRF and ALF mice. E. Metabolite set enrichment analysis (MSEA) of the deregulated metabolites. Size of the dot represents the enrichment ratio. F. Pairwise correlation of significantly deregulated (p-value ≤ 0.05; log_2_ Fold change ALF/TRF≥±1.0) proteins and metabolites in the liver. Statistical significance was determined using unpaired t-test with Welch’s correction in A. Each dot represents a biological replicate.
3

### Slide 4
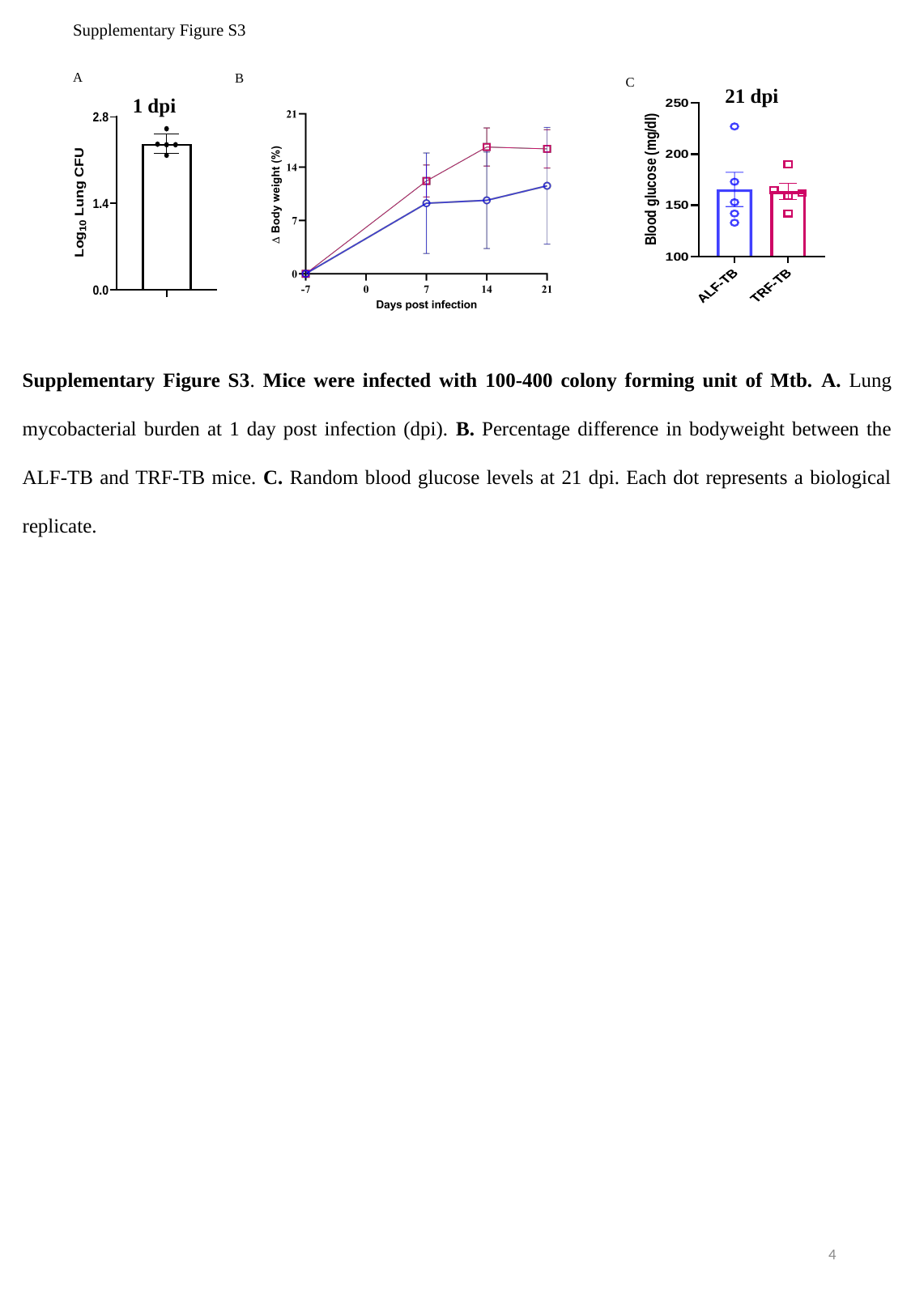

Supplementary Figure S3
A
B
C
21 dpi
1 dpi
Supplementary Figure S3. Mice were infected with 100-400 colony forming unit of Mtb. A. Lung mycobacterial burden at 1 day post infection (dpi). B. Percentage difference in bodyweight between the ALF-TB and TRF-TB mice. C. Random blood glucose levels at 21 dpi. Each dot represents a biological replicate.
4

### Slide 5
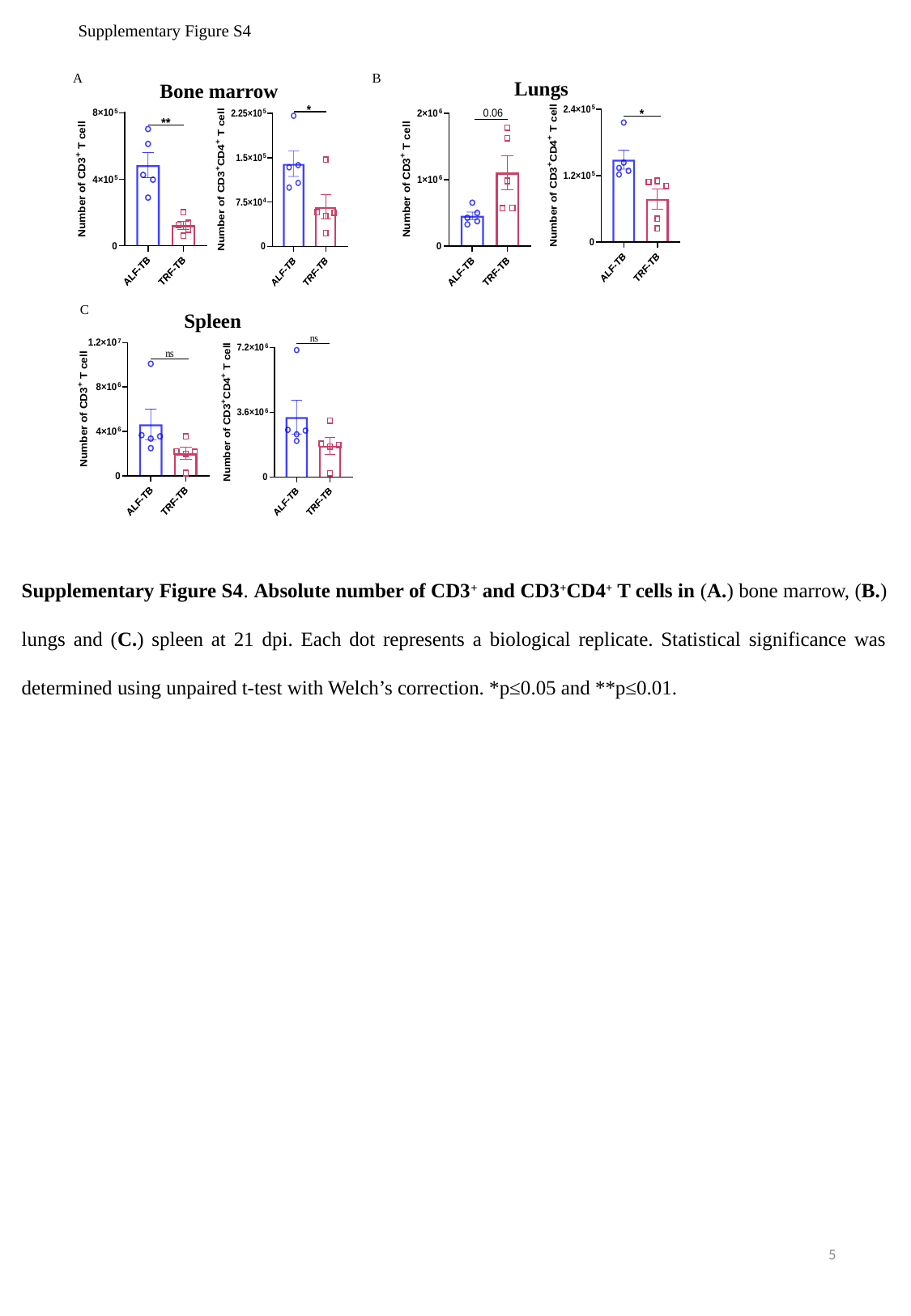

Supplementary Figure S4
A
B
Lungs
Bone marrow
C
Spleen
Supplementary Figure S4. Absolute number of CD3^+^ and CD3^+^CD4^+^ T cells in (A.) bone marrow, (B.) lungs and (C.) spleen at 21 dpi. Each dot represents a biological replicate. Statistical significance was determined using unpaired t-test with Welch’s correction. *p≤0.05 and **p≤0.01.
5

### Slide 6
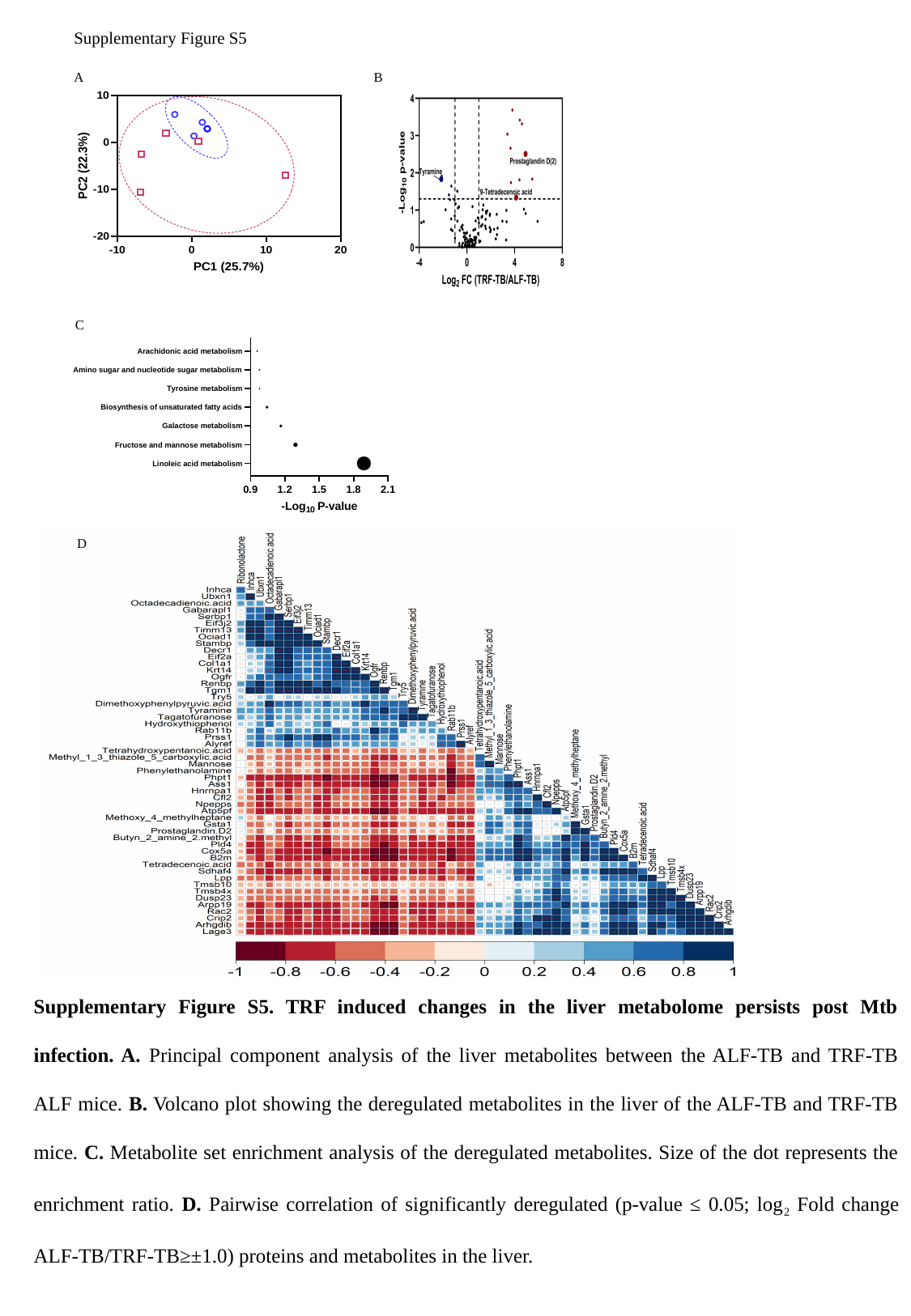

Supplementary Figure S5
A
B
C
D
Supplementary Figure S5. TRF induced changes in the liver metabolome persists post Mtb infection. A. Principal component analysis of the liver metabolites between the ALF-TB and TRF-TB ALF mice. B. Volcano plot showing the deregulated metabolites in the liver of the ALF-TB and TRF-TB mice. C. Metabolite set enrichment analysis of the deregulated metabolites. Size of the dot represents the enrichment ratio. D. Pairwise correlation of significantly deregulated (p-value ≤ 0.05; log_2_ Fold change ALF-TB/TRF-TB≥±1.0) proteins and metabolites in the liver.

### Slide 7
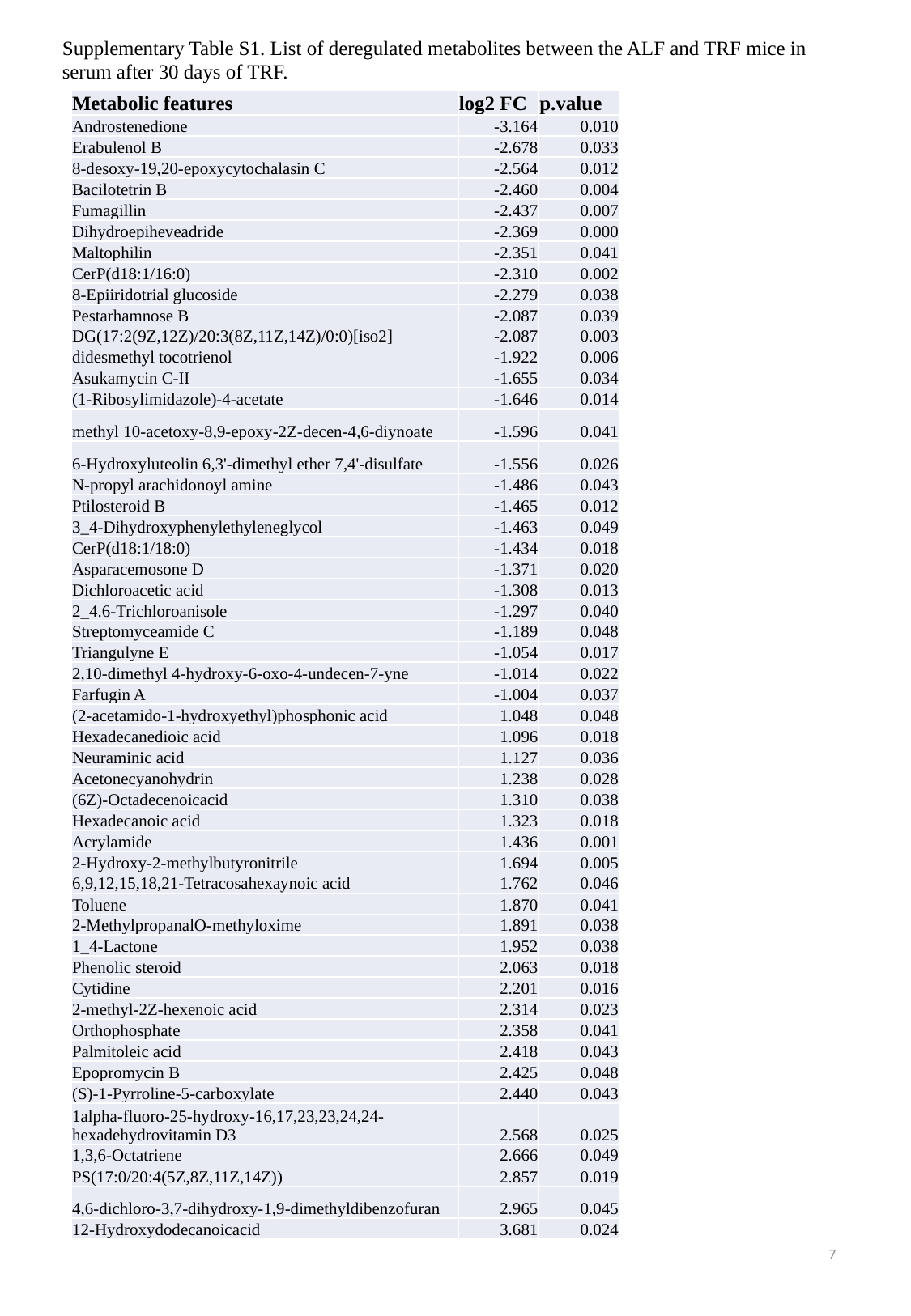

Supplementary Table S1. List of deregulated metabolites between the ALF and TRF mice in serum after 30 days of TRF.
| Metabolic features | log2 FC | p.value |
| --- | --- | --- |
| Androstenedione | -3.164 | 0.010 |
| Erabulenol B | -2.678 | 0.033 |
| 8-desoxy-19,20-epoxycytochalasin C | -2.564 | 0.012 |
| Bacilotetrin B | -2.460 | 0.004 |
| Fumagillin | -2.437 | 0.007 |
| Dihydroepiheveadride | -2.369 | 0.000 |
| Maltophilin | -2.351 | 0.041 |
| CerP(d18:1/16:0) | -2.310 | 0.002 |
| 8-Epiiridotrial glucoside | -2.279 | 0.038 |
| Pestarhamnose B | -2.087 | 0.039 |
| DG(17:2(9Z,12Z)/20:3(8Z,11Z,14Z)/0:0)[iso2] | -2.087 | 0.003 |
| didesmethyl tocotrienol | -1.922 | 0.006 |
| Asukamycin C-II | -1.655 | 0.034 |
| (1-Ribosylimidazole)-4-acetate | -1.646 | 0.014 |
| methyl 10-acetoxy-8,9-epoxy-2Z-decen-4,6-diynoate | -1.596 | 0.041 |
| 6-Hydroxyluteolin 6,3'-dimethyl ether 7,4'-disulfate | -1.556 | 0.026 |
| N-propyl arachidonoyl amine | -1.486 | 0.043 |
| Ptilosteroid B | -1.465 | 0.012 |
| 3\_4-Dihydroxyphenylethyleneglycol | -1.463 | 0.049 |
| CerP(d18:1/18:0) | -1.434 | 0.018 |
| Asparacemosone D | -1.371 | 0.020 |
| Dichloroacetic acid | -1.308 | 0.013 |
| 2\_4.6-Trichloroanisole | -1.297 | 0.040 |
| Streptomyceamide C | -1.189 | 0.048 |
| Triangulyne E | -1.054 | 0.017 |
| 2,10-dimethyl 4-hydroxy-6-oxo-4-undecen-7-yne | -1.014 | 0.022 |
| Farfugin A | -1.004 | 0.037 |
| (2-acetamido-1-hydroxyethyl)phosphonic acid | 1.048 | 0.048 |
| Hexadecanedioic acid | 1.096 | 0.018 |
| Neuraminic acid | 1.127 | 0.036 |
| Acetonecyanohydrin | 1.238 | 0.028 |
| (6Z)-Octadecenoicacid | 1.310 | 0.038 |
| Hexadecanoic acid | 1.323 | 0.018 |
| Acrylamide | 1.436 | 0.001 |
| 2-Hydroxy-2-methylbutyronitrile | 1.694 | 0.005 |
| 6,9,12,15,18,21-Tetracosahexaynoic acid | 1.762 | 0.046 |
| Toluene | 1.870 | 0.041 |
| 2-MethylpropanalO-methyloxime | 1.891 | 0.038 |
| 1\_4-Lactone | 1.952 | 0.038 |
| Phenolic steroid | 2.063 | 0.018 |
| Cytidine | 2.201 | 0.016 |
| 2-methyl-2Z-hexenoic acid | 2.314 | 0.023 |
| Orthophosphate | 2.358 | 0.041 |
| Palmitoleic acid | 2.418 | 0.043 |
| Epopromycin B | 2.425 | 0.048 |
| (S)-1-Pyrroline-5-carboxylate | 2.440 | 0.043 |
| 1alpha-fluoro-25-hydroxy-16,17,23,23,24,24-hexadehydrovitamin D3 | 2.568 | 0.025 |
| 1,3,6-Octatriene | 2.666 | 0.049 |
| PS(17:0/20:4(5Z,8Z,11Z,14Z)) | 2.857 | 0.019 |
| 4,6-dichloro-3,7-dihydroxy-1,9-dimethyldibenzofuran | 2.965 | 0.045 |
| 12-Hydroxydodecanoicacid | 3.681 | 0.024 |
7

### Slide 8
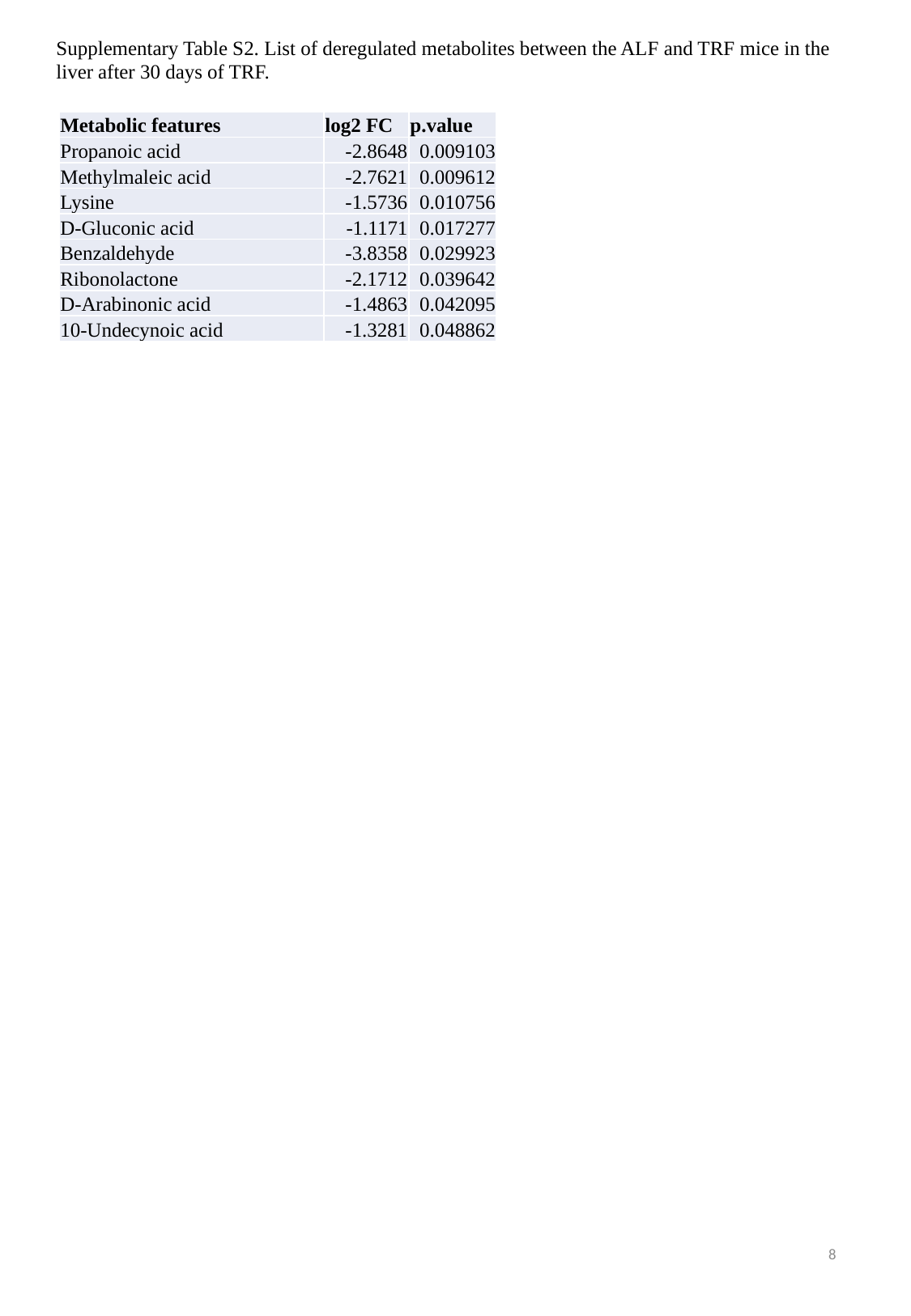

Supplementary Table S2. List of deregulated metabolites between the ALF and TRF mice in the liver after 30 days of TRF.
| Metabolic features | log2 FC | p.value |
| --- | --- | --- |
| Propanoic acid | -2.8648 | 0.009103 |
| Methylmaleic acid | -2.7621 | 0.009612 |
| Lysine | -1.5736 | 0.010756 |
| D-Gluconic acid | -1.1171 | 0.017277 |
| Benzaldehyde | -3.8358 | 0.029923 |
| Ribonolactone | -2.1712 | 0.039642 |
| D-Arabinonic acid | -1.4863 | 0.042095 |
| 10-Undecynoic acid | -1.3281 | 0.048862 |
8

### Slide 9
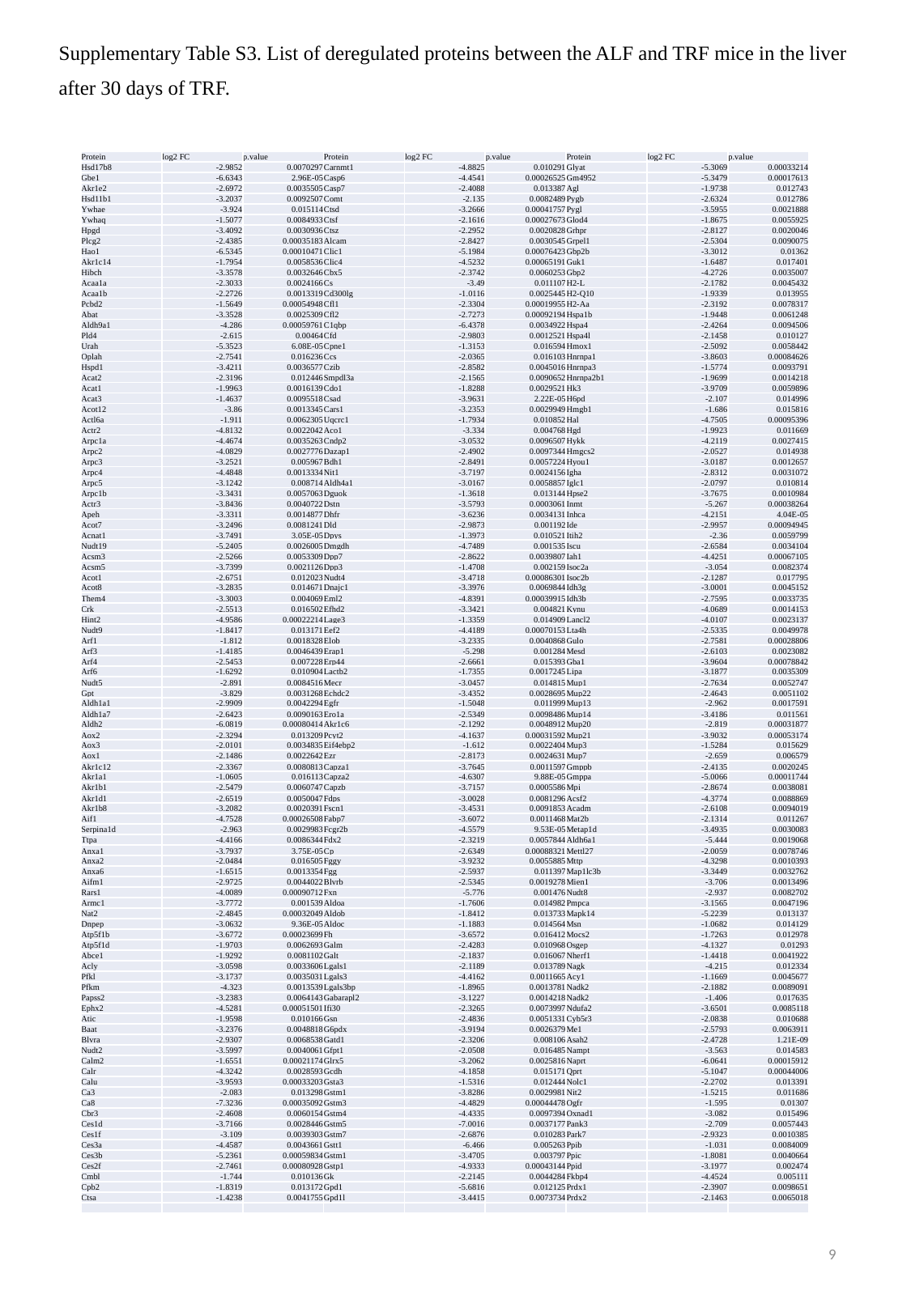

Supplementary Table S3. List of deregulated proteins between the ALF and TRF mice in the liver after 30 days of TRF.
| Protein | log2 FC | p.value | Protein | log2 FC | p.value | Protein | log2 FC | p.value |
| --- | --- | --- | --- | --- | --- | --- | --- | --- |
| Hsd17b8 | -2.9852 | 0.0070297 | Carnmt1 | -4.8825 | 0.010291 | Glyat | -5.3069 | 0.00033214 |
| Gbe1 | -6.6343 | 2.96E-05 | Casp6 | -4.4541 | 0.00026525 | Gm4952 | -5.3479 | 0.00017613 |
| Akr1e2 | -2.6972 | 0.0035505 | Casp7 | -2.4088 | 0.013387 | Agl | -1.9738 | 0.012743 |
| Hsd11b1 | -3.2037 | 0.0092507 | Comt | -2.135 | 0.0082489 | Pygb | -2.6324 | 0.012786 |
| Ywhae | -3.924 | 0.015114 | Ctsd | -3.2666 | 0.00041757 | Pygl | -3.5955 | 0.0021888 |
| Ywhaq | -1.5077 | 0.0084933 | Ctsf | -2.1616 | 0.00027673 | Glod4 | -1.8675 | 0.0055925 |
| Hpgd | -3.4092 | 0.0030936 | Ctsz | -2.2952 | 0.0020828 | Grhpr | -2.8127 | 0.0020046 |
| Plcg2 | -2.4385 | 0.00035183 | Alcam | -2.8427 | 0.0030545 | Grpel1 | -2.5304 | 0.0090075 |
| Hao1 | -6.5345 | 0.00010471 | Clic1 | -5.1984 | 0.00076423 | Gbp2b | -3.3012 | 0.01362 |
| Akr1c14 | -1.7954 | 0.0058536 | Clic4 | -4.5232 | 0.00065191 | Guk1 | -1.6487 | 0.017401 |
| Hibch | -3.3578 | 0.0032646 | Cbx5 | -2.3742 | 0.0060253 | Gbp2 | -4.2726 | 0.0035007 |
| Acaa1a | -2.3033 | 0.0024166 | Cs | -3.49 | 0.011107 | H2-L | -2.1782 | 0.0045432 |
| Acaa1b | -2.2726 | 0.0013319 | Cd300lg | -1.0116 | 0.0025445 | H2-Q10 | -1.9339 | 0.013955 |
| Pcbd2 | -1.5649 | 0.00054948 | Cfl1 | -2.3304 | 0.00019955 | H2-Aa | -2.3192 | 0.0078317 |
| Abat | -3.3528 | 0.0025309 | Cfl2 | -2.7273 | 0.00092194 | Hspa1b | -1.9448 | 0.0061248 |
| Aldh9a1 | -4.286 | 0.00059761 | C1qbp | -6.4378 | 0.0034922 | Hspa4 | -2.4264 | 0.0094506 |
| Pld4 | -2.615 | 0.00464 | Cfd | -2.9803 | 0.0012521 | Hspa4l | -2.1458 | 0.010127 |
| Urah | -5.3523 | 6.08E-05 | Cpne1 | -1.3153 | 0.016594 | Hmox1 | -2.5092 | 0.0058442 |
| Oplah | -2.7541 | 0.016236 | Ccs | -2.0365 | 0.016103 | Hnrnpa1 | -3.8603 | 0.00084626 |
| Hspd1 | -3.4211 | 0.0036577 | Czib | -2.8582 | 0.0045016 | Hnrnpa3 | -1.5774 | 0.0093791 |
| Acat2 | -2.3196 | 0.012446 | Smpdl3a | -2.1565 | 0.0090652 | Hnrnpa2b1 | -1.9699 | 0.0014218 |
| Acat1 | -1.9963 | 0.0016139 | Cdo1 | -1.8288 | 0.0029521 | Hk3 | -3.9709 | 0.0059896 |
| Acat3 | -1.4637 | 0.0095518 | Csad | -3.9631 | 2.22E-05 | H6pd | -2.107 | 0.014996 |
| Acot12 | -3.86 | 0.0013345 | Cars1 | -3.2353 | 0.0029949 | Hmgb1 | -1.686 | 0.015816 |
| Actl6a | -1.911 | 0.0062305 | Uqcrc1 | -1.7934 | 0.010852 | Hal | -4.7505 | 0.00095396 |
| Actr2 | -4.8132 | 0.0022042 | Aco1 | -3.334 | 0.004768 | Hgd | -1.9923 | 0.011669 |
| Arpc1a | -4.4674 | 0.0035263 | Cndp2 | -3.0532 | 0.0096507 | Hykk | -4.2119 | 0.0027415 |
| Arpc2 | -4.0829 | 0.0027776 | Dazap1 | -2.4902 | 0.0097344 | Hmgcs2 | -2.0527 | 0.014938 |
| Arpc3 | -3.2521 | 0.005967 | Bdh1 | -2.8491 | 0.0057224 | Hyou1 | -3.0187 | 0.0012657 |
| Arpc4 | -4.4848 | 0.0013334 | Nit1 | -3.7197 | 0.0024156 | Igha | -2.8312 | 0.0031072 |
| Arpc5 | -3.1242 | 0.008714 | Aldh4a1 | -3.0167 | 0.0058857 | Iglc1 | -2.0797 | 0.010814 |
| Arpc1b | -3.3431 | 0.0057063 | Dguok | -1.3618 | 0.013144 | Hpse2 | -3.7675 | 0.0010984 |
| Actr3 | -3.8436 | 0.0040722 | Dstn | -3.5793 | 0.0003061 | Inmt | -5.267 | 0.00038264 |
| Apeh | -3.3311 | 0.0014877 | Dhfr | -3.6236 | 0.0034131 | Inhca | -4.2151 | 4.04E-05 |
| Acot7 | -3.2496 | 0.0081241 | Dld | -2.9873 | 0.001192 | Ide | -2.9957 | 0.00094945 |
| Acnat1 | -3.7491 | 3.05E-05 | Dpys | -1.3973 | 0.010521 | Itih2 | -2.36 | 0.0059799 |
| Nudt19 | -5.2405 | 0.0026005 | Dmgdh | -4.7489 | 0.001535 | Iscu | -2.6584 | 0.0034104 |
| Acsm3 | -2.5266 | 0.0053309 | Dpp7 | -2.8622 | 0.0039807 | Iah1 | -4.4251 | 0.00067105 |
| Acsm5 | -3.7399 | 0.0021126 | Dpp3 | -1.4708 | 0.002159 | Isoc2a | -3.054 | 0.0082374 |
| Acot1 | -2.6751 | 0.012023 | Nudt4 | -3.4718 | 0.00086301 | Isoc2b | -2.1287 | 0.017795 |
| Acot8 | -3.2835 | 0.014671 | Dnajc1 | -3.3976 | 0.0069844 | Idh3g | -3.0001 | 0.0045152 |
| Them4 | -3.3003 | 0.004069 | Eml2 | -4.8391 | 0.00039915 | Idh3b | -2.7595 | 0.0033735 |
| Crk | -2.5513 | 0.016502 | Efhd2 | -3.3421 | 0.004821 | Kynu | -4.0689 | 0.0014153 |
| Hint2 | -4.9586 | 0.00022214 | Lage3 | -1.3359 | 0.014909 | Lancl2 | -4.0107 | 0.0023137 |
| Nudt9 | -1.8417 | 0.013171 | Eef2 | -4.4189 | 0.00070153 | Lta4h | -2.5335 | 0.0049978 |
| Arf1 | -1.812 | 0.0018328 | Elob | -3.2335 | 0.0040868 | Gulo | -2.7581 | 0.00028806 |
| Arf3 | -1.4185 | 0.0046439 | Erap1 | -5.298 | 0.001284 | Mesd | -2.6103 | 0.0023082 |
| Arf4 | -2.5453 | 0.007228 | Erp44 | -2.6661 | 0.015393 | Gba1 | -3.9604 | 0.00078842 |
| Arf6 | -1.6292 | 0.010904 | Lactb2 | -1.7355 | 0.0017245 | Lipa | -3.1877 | 0.0035309 |
| Nudt5 | -2.891 | 0.0084516 | Mecr | -3.0457 | 0.014815 | Mup1 | -2.7634 | 0.0052747 |
| Gpt | -3.829 | 0.0031268 | Echdc2 | -3.4352 | 0.0028695 | Mup22 | -2.4643 | 0.0051102 |
| Aldh1a1 | -2.9909 | 0.0042294 | Egfr | -1.5048 | 0.011999 | Mup13 | -2.962 | 0.0017591 |
| Aldh1a7 | -2.6423 | 0.0090163 | Ero1a | -2.5349 | 0.0098486 | Mup14 | -3.4186 | 0.011561 |
| Aldh2 | -6.0819 | 0.00080414 | Akr1c6 | -2.1292 | 0.0048912 | Mup20 | -2.819 | 0.00031877 |
| Aox2 | -2.3294 | 0.013209 | Pcyt2 | -4.1637 | 0.00031592 | Mup21 | -3.9032 | 0.00053174 |
| Aox3 | -2.0101 | 0.0034835 | Eif4ebp2 | -1.612 | 0.0022404 | Mup3 | -1.5284 | 0.015629 |
| Aox1 | -2.1486 | 0.0022642 | Ezr | -2.8173 | 0.0024631 | Mup7 | -2.659 | 0.006579 |
| Akr1c12 | -2.3367 | 0.0080813 | Capza1 | -3.7645 | 0.0011597 | Gmppb | -2.4135 | 0.0020245 |
| Akr1a1 | -1.0605 | 0.016113 | Capza2 | -4.6307 | 9.88E-05 | Gmppa | -5.0066 | 0.00011744 |
| Akr1b1 | -2.5479 | 0.0060747 | Capzb | -3.7157 | 0.0005586 | Mpi | -2.8674 | 0.0038081 |
| Akr1d1 | -2.6519 | 0.0050047 | Fdps | -3.0028 | 0.0081296 | Acsf2 | -4.3774 | 0.0088869 |
| Akr1b8 | -3.2082 | 0.0020391 | Fscn1 | -3.4531 | 0.0091853 | Acadm | -2.6108 | 0.0094019 |
| Aif1 | -4.7528 | 0.00026508 | Fabp7 | -3.6072 | 0.0011468 | Mat2b | -2.1314 | 0.011267 |
| Serpina1d | -2.963 | 0.0029983 | Fcgr2b | -4.5579 | 9.53E-05 | Metap1d | -3.4935 | 0.0030083 |
| Ttpa | -4.4166 | 0.0086344 | Fdx2 | -2.3219 | 0.0057844 | Aldh6a1 | -5.444 | 0.0019068 |
| Anxa1 | -3.7937 | 3.75E-05 | Cp | -2.6349 | 0.00088321 | Mettl27 | -2.0059 | 0.0078746 |
| Anxa2 | -2.0484 | 0.016505 | Fggy | -3.9232 | 0.0055885 | Mttp | -4.3298 | 0.0010393 |
| Anxa6 | -1.6515 | 0.0013354 | Fgg | -2.5937 | 0.011397 | Map1lc3b | -3.3449 | 0.0032762 |
| Aifm1 | -2.9725 | 0.0044022 | Blvrb | -2.5345 | 0.0019278 | Mien1 | -3.706 | 0.0013496 |
| Rars1 | -4.0089 | 0.00090712 | Fxn | -5.776 | 0.001476 | Nudt8 | -2.937 | 0.0082702 |
| Armc1 | -3.7772 | 0.001539 | Aldoa | -1.7606 | 0.014982 | Pmpca | -3.1565 | 0.0047196 |
| Nat2 | -2.4845 | 0.00032049 | Aldob | -1.8412 | 0.013733 | Mapk14 | -5.2239 | 0.013137 |
| Dnpep | -3.0632 | 9.36E-05 | Aldoc | -1.1883 | 0.014564 | Msn | -1.0682 | 0.014129 |
| Atp5f1b | -3.6772 | 0.00023699 | Fh | -3.6572 | 0.016412 | Mocs2 | -1.7263 | 0.012978 |
| Atp5f1d | -1.9703 | 0.0062693 | Galm | -2.4283 | 0.010968 | Osgep | -4.1327 | 0.01293 |
| Abce1 | -1.9292 | 0.0081102 | Galt | -2.1837 | 0.016067 | Nherf1 | -1.4418 | 0.0041922 |
| Acly | -3.0598 | 0.0033606 | Lgals1 | -2.1189 | 0.013789 | Nagk | -4.215 | 0.012334 |
| Pfkl | -3.1737 | 0.0035031 | Lgals3 | -4.4162 | 0.0011665 | Acy1 | -1.1669 | 0.0045677 |
| Pfkm | -4.323 | 0.0013539 | Lgals3bp | -1.8965 | 0.0013781 | Nadk2 | -2.1882 | 0.0089091 |
| Papss2 | -3.2383 | 0.0064143 | Gabarapl2 | -3.1227 | 0.0014218 | Nadk2 | -1.406 | 0.017635 |
| Ephx2 | -4.5281 | 0.00051501 | Ifi30 | -2.3265 | 0.0073997 | Ndufa2 | -3.6501 | 0.0085118 |
| Atic | -1.9598 | 0.010166 | Gsn | -2.4836 | 0.0051331 | Cyb5r3 | -2.0838 | 0.010688 |
| Baat | -3.2376 | 0.0048818 | G6pdx | -3.9194 | 0.0026379 | Me1 | -2.5793 | 0.0063911 |
| Blvra | -2.9307 | 0.0068538 | Gatd1 | -2.3206 | 0.008106 | Asah2 | -2.4728 | 1.21E-09 |
| Nudt2 | -3.5997 | 0.0040061 | Gfpt1 | -2.0508 | 0.016485 | Nampt | -3.563 | 0.014583 |
| Calm2 | -1.6551 | 0.00021174 | Glrx5 | -3.2062 | 0.0025816 | Naprt | -6.0641 | 0.00015912 |
| Calr | -4.3242 | 0.0028593 | Gcdh | -4.1858 | 0.015171 | Qprt | -5.1047 | 0.00044006 |
| Calu | -3.9593 | 0.00033203 | Gsta3 | -1.5316 | 0.012444 | Nolc1 | -2.2702 | 0.013391 |
| Ca3 | -2.083 | 0.013298 | Gstm1 | -3.8286 | 0.0029981 | Nit2 | -1.5215 | 0.011686 |
| Ca8 | -7.3236 | 0.00035092 | Gstm3 | -4.4829 | 0.00044478 | Ogfr | -1.595 | 0.01307 |
| Cbr3 | -2.4608 | 0.0060154 | Gstm4 | -4.4335 | 0.0097394 | Oxnad1 | -3.082 | 0.015496 |
| Ces1d | -3.7166 | 0.0028446 | Gstm5 | -7.0016 | 0.0037177 | Pank3 | -2.709 | 0.0057443 |
| Ces1f | -3.109 | 0.0039303 | Gstm7 | -2.6876 | 0.010283 | Park7 | -2.9323 | 0.0010385 |
| Ces3a | -4.4587 | 0.0043661 | Gstt1 | -6.466 | 0.005263 | Ppib | -1.031 | 0.0084009 |
| Ces3b | -5.2361 | 0.00059834 | Gstm1 | -3.4705 | 0.003797 | Ppic | -1.8081 | 0.0040664 |
| Ces2f | -2.7461 | 0.00080928 | Gstp1 | -4.9333 | 0.00043144 | Ppid | -3.1977 | 0.002474 |
| Cmbl | -1.744 | 0.010136 | Gk | -2.2145 | 0.0044284 | Fkbp4 | -4.4524 | 0.005111 |
| Cpb2 | -1.8319 | 0.013172 | Gpd1 | -5.6816 | 0.012125 | Prdx1 | -2.3907 | 0.0098651 |
| Ctsa | -1.4238 | 0.0041755 | Gpd1l | -3.4415 | 0.0073734 | Prdx2 | -2.1463 | 0.0065018 |
9

### Slide 10
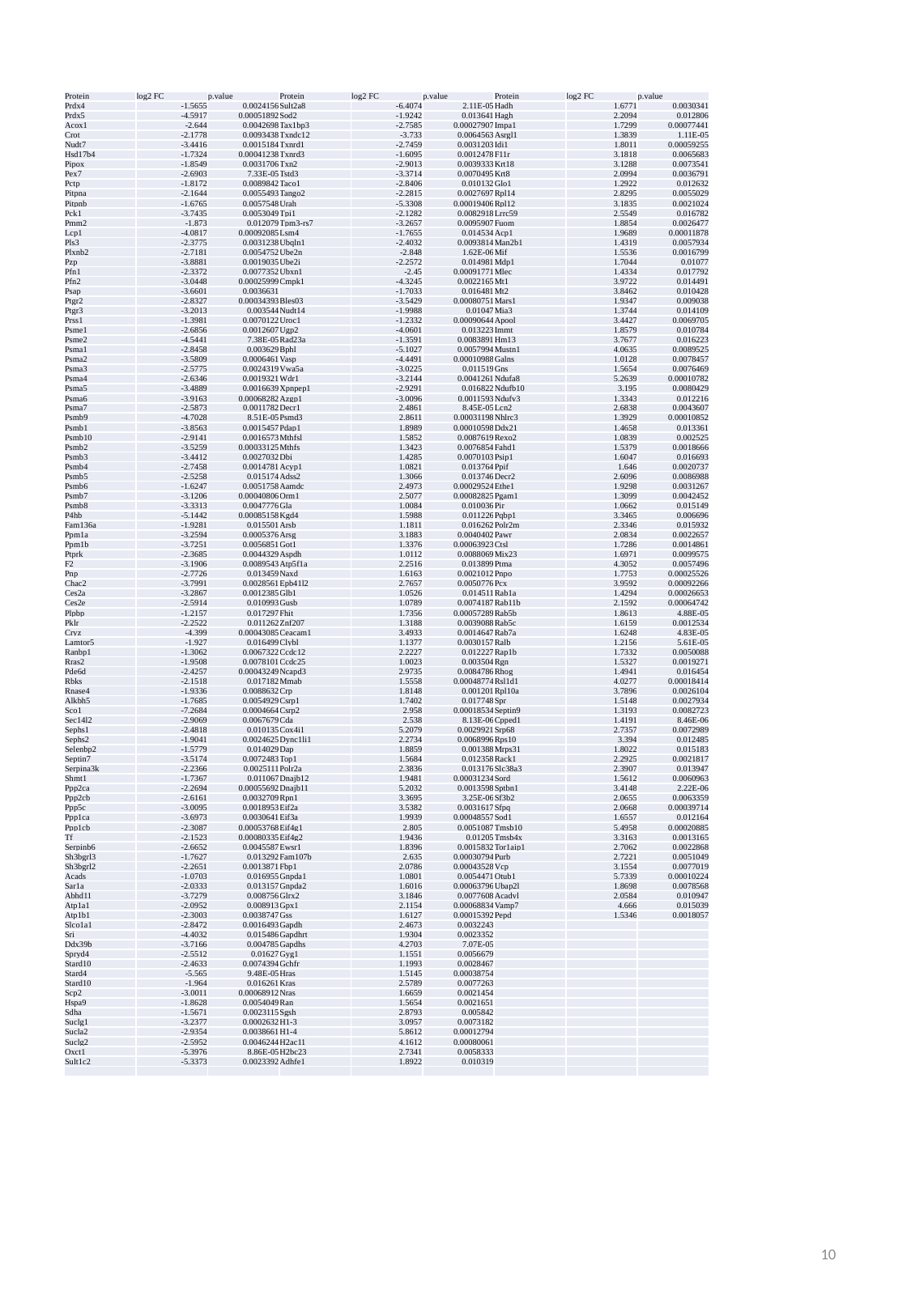

| Protein | log2 FC | p.value | Protein | log2 FC | p.value | Protein | log2 FC | p.value |
| --- | --- | --- | --- | --- | --- | --- | --- | --- |
| Prdx4 | -1.5655 | 0.0024156 | Sult2a8 | -6.4074 | 2.11E-05 | Hadh | 1.6771 | 0.0030341 |
| Prdx5 | -4.5917 | 0.00051892 | Sod2 | -1.9242 | 0.013641 | Hagh | 2.2094 | 0.012806 |
| Acox1 | -2.644 | 0.0042698 | Tax1bp3 | -2.7585 | 0.00027907 | Impa1 | 1.7299 | 0.00077441 |
| Crot | -2.1778 | 0.0093438 | Txndc12 | -3.733 | 0.0064563 | Asrgl1 | 1.3839 | 1.11E-05 |
| Nudt7 | -3.4416 | 0.0015184 | Txnrd1 | -2.7459 | 0.0031203 | Idi1 | 1.8011 | 0.00059255 |
| Hsd17b4 | -1.7324 | 0.00041238 | Txnrd3 | -1.6095 | 0.0012478 | F11r | 3.1818 | 0.0065683 |
| Pipox | -1.8549 | 0.0031706 | Txn2 | -2.9013 | 0.0039333 | Krt18 | 3.1288 | 0.0073541 |
| Pex7 | -2.6903 | 7.33E-05 | Tstd3 | -3.3714 | 0.0070495 | Krt8 | 2.0994 | 0.0036791 |
| Pctp | -1.8172 | 0.0089842 | Taco1 | -2.8406 | 0.010132 | Glo1 | 1.2922 | 0.012632 |
| Pitpna | -2.1644 | 0.0055493 | Tango2 | -2.2815 | 0.0027697 | Rpl14 | 2.8295 | 0.0055029 |
| Pitpnb | -1.6765 | 0.0057548 | Urah | -5.3308 | 0.00019406 | Rpl12 | 3.1835 | 0.0021024 |
| Pck1 | -3.7435 | 0.0053049 | Tpi1 | -2.1282 | 0.0082918 | Lrrc59 | 2.5549 | 0.016782 |
| Pmm2 | -1.873 | 0.012079 | Tpm3-rs7 | -3.2657 | 0.0095907 | Fuom | 1.8854 | 0.0026477 |
| Lcp1 | -4.0817 | 0.00092085 | Lsm4 | -1.7655 | 0.014534 | Acp1 | 1.9689 | 0.00011878 |
| Pls3 | -2.3775 | 0.0031238 | Ubqln1 | -2.4032 | 0.0093814 | Man2b1 | 1.4319 | 0.0057934 |
| Plxnb2 | -2.7181 | 0.0054752 | Ube2n | -2.848 | 1.62E-06 | Mif | 1.5536 | 0.0016799 |
| Pzp | -3.8881 | 0.0019035 | Ube2i | -2.2572 | 0.014981 | Mdp1 | 1.7044 | 0.01077 |
| Pfn1 | -2.3372 | 0.0077352 | Ubxn1 | -2.45 | 0.00091771 | Mlec | 1.4334 | 0.017792 |
| Pfn2 | -3.0448 | 0.00025999 | Cmpk1 | -4.3245 | 0.0022165 | Mt1 | 3.9722 | 0.014491 |
| Psap | -3.6601 | 0.0036631 | | -1.7033 | 0.016481 | Mt2 | 3.8462 | 0.010428 |
| Ptgr2 | -2.8327 | 0.00034393 | Bles03 | -3.5429 | 0.00080751 | Mars1 | 1.9347 | 0.009038 |
| Ptgr3 | -3.2013 | 0.003544 | Nudt14 | -1.9988 | 0.01047 | Mia3 | 1.3744 | 0.014109 |
| Prss1 | -1.3981 | 0.0070122 | Uroc1 | -1.2332 | 0.00090644 | Apool | 3.4427 | 0.0069705 |
| Psme1 | -2.6856 | 0.0012607 | Ugp2 | -4.0601 | 0.013223 | Immt | 1.8579 | 0.010784 |
| Psme2 | -4.5441 | 7.38E-05 | Rad23a | -1.3591 | 0.0083891 | Hm13 | 3.7677 | 0.016223 |
| Psma1 | -2.8458 | 0.003629 | Bphl | -5.1027 | 0.0057994 | Mustn1 | 4.0635 | 0.0089525 |
| Psma2 | -3.5809 | 0.0006461 | Vasp | -4.4491 | 0.00010988 | Galns | 1.0128 | 0.0078457 |
| Psma3 | -2.5775 | 0.0024319 | Vwa5a | -3.0225 | 0.011519 | Gns | 1.5654 | 0.0076469 |
| Psma4 | -2.6346 | 0.0019321 | Wdr1 | -3.2144 | 0.0041261 | Ndufa8 | 5.2639 | 0.00010782 |
| Psma5 | -3.4889 | 0.0016639 | Xpnpep1 | -2.9291 | 0.016822 | Ndufb10 | 3.195 | 0.0080429 |
| Psma6 | -3.9163 | 0.00068282 | Azgp1 | -3.0096 | 0.0011593 | Ndufv3 | 1.3343 | 0.012216 |
| Psma7 | -2.5873 | 0.0011782 | Decr1 | 2.4861 | 8.45E-05 | Lcn2 | 2.6838 | 0.0043607 |
| Psmb9 | -4.7028 | 8.51E-05 | Psmd3 | 2.8611 | 0.00031198 | Nhlrc3 | 1.3929 | 0.00010852 |
| Psmb1 | -3.8563 | 0.0015457 | Pdap1 | 1.8989 | 0.00010598 | Ddx21 | 1.4658 | 0.013361 |
| Psmb10 | -2.9141 | 0.0016573 | Mthfsl | 1.5852 | 0.0087619 | Rexo2 | 1.0839 | 0.002525 |
| Psmb2 | -3.5259 | 0.00033125 | Mthfs | 1.3423 | 0.0076854 | Fahd1 | 1.5379 | 0.0018666 |
| Psmb3 | -3.4412 | 0.0027032 | Dbi | 1.4285 | 0.0070103 | Psip1 | 1.6047 | 0.016693 |
| Psmb4 | -2.7458 | 0.0014781 | Acyp1 | 1.0821 | 0.013764 | Ppif | 1.646 | 0.0020737 |
| Psmb5 | -2.5258 | 0.015174 | Adss2 | 1.3066 | 0.013746 | Decr2 | 2.6096 | 0.0086988 |
| Psmb6 | -1.6247 | 0.0051758 | Aamdc | 2.4973 | 0.00029524 | Ethe1 | 1.9298 | 0.0031267 |
| Psmb7 | -3.1206 | 0.00040806 | Orm1 | 2.5077 | 0.00082825 | Pgam1 | 1.3099 | 0.0042452 |
| Psmb8 | -3.3313 | 0.0047776 | Gla | 1.0084 | 0.010036 | Pir | 1.0662 | 0.015149 |
| P4hb | -5.1442 | 0.00085158 | Kgd4 | 1.5988 | 0.011226 | Pqbp1 | 3.3465 | 0.006696 |
| Fam136a | -1.9281 | 0.015501 | Arsb | 1.1811 | 0.016262 | Polr2m | 2.3346 | 0.015932 |
| Ppm1a | -3.2594 | 0.0005376 | Arsg | 3.1883 | 0.0040402 | Pawr | 2.0834 | 0.0022657 |
| Ppm1b | -3.7251 | 0.0056851 | Got1 | 1.3376 | 0.00063923 | Ctsl | 1.7286 | 0.0014861 |
| Ptprk | -2.3685 | 0.0044329 | Aspdh | 1.0112 | 0.0088069 | Mix23 | 1.6971 | 0.0099575 |
| F2 | -3.1906 | 0.0089543 | Atp5f1a | 2.2516 | 0.013899 | Ptma | 4.3052 | 0.0057496 |
| Pnp | -2.7726 | 0.013459 | Naxd | 1.6163 | 0.0021012 | Pnpo | 1.7753 | 0.00025526 |
| Chac2 | -3.7991 | 0.0028561 | Epb41l2 | 2.7657 | 0.0050776 | Pcx | 3.9592 | 0.00092266 |
| Ces2a | -3.2867 | 0.0012385 | Glb1 | 1.0526 | 0.014511 | Rab1a | 1.4294 | 0.00026653 |
| Ces2e | -2.5914 | 0.010993 | Gusb | 1.0789 | 0.0074187 | Rab11b | 2.1592 | 0.00064742 |
| Plpbp | -1.2157 | 0.017297 | Fhit | 1.7356 | 0.00057289 | Rab5b | 1.8613 | 4.88E-05 |
| Pklr | -2.2522 | 0.011262 | Znf207 | 1.3188 | 0.0039088 | Rab5c | 1.6159 | 0.0012534 |
| Cryz | -4.399 | 0.00043085 | Ceacam1 | 3.4933 | 0.0014647 | Rab7a | 1.6248 | 4.83E-05 |
| Lamtor5 | -1.927 | 0.016499 | Clybl | 1.1377 | 0.0030157 | Ralb | 1.2156 | 5.61E-05 |
| Ranbp1 | -1.3062 | 0.0067322 | Ccdc12 | 2.2227 | 0.012227 | Rap1b | 1.7332 | 0.0050088 |
| Rras2 | -1.9508 | 0.0078101 | Ccdc25 | 1.0023 | 0.003504 | Rgn | 1.5327 | 0.0019271 |
| Pde6d | -2.4257 | 0.00043249 | Ncapd3 | 2.9735 | 0.0084786 | Rhog | 1.4941 | 0.016454 |
| Rbks | -2.1518 | 0.017182 | Mmab | 1.5558 | 0.00048774 | Rsl1d1 | 4.0277 | 0.00018414 |
| Rnase4 | -1.9336 | 0.0088632 | Crp | 1.8148 | 0.001201 | Rpl10a | 3.7896 | 0.0026104 |
| Alkbh5 | -1.7685 | 0.0054929 | Csrp1 | 1.7402 | 0.017748 | Spr | 1.5148 | 0.0027934 |
| Sco1 | -7.2684 | 0.0004664 | Csrp2 | 2.958 | 0.00018534 | Septin9 | 1.3193 | 0.0082723 |
| Sec14l2 | -2.9069 | 0.0067679 | Cda | 2.538 | 8.13E-06 | Cpped1 | 1.4191 | 8.46E-06 |
| Sephs1 | -2.4818 | 0.010135 | Cox4i1 | 5.2079 | 0.0029921 | Srp68 | 2.7357 | 0.0072989 |
| Sephs2 | -1.9041 | 0.0024625 | Dync1li1 | 2.2734 | 0.0068996 | Rps10 | 3.394 | 0.012485 |
| Selenbp2 | -1.5779 | 0.014029 | Dap | 1.8859 | 0.001388 | Mrps31 | 1.8022 | 0.015183 |
| Septin7 | -3.5174 | 0.0072483 | Top1 | 1.5684 | 0.012358 | Rack1 | 2.2925 | 0.0021817 |
| Serpina3k | -2.2366 | 0.0025111 | Polr2a | 2.3836 | 0.013176 | Slc38a3 | 2.3907 | 0.013947 |
| Shmt1 | -1.7367 | 0.011067 | Dnajb12 | 1.9481 | 0.00031234 | Sord | 1.5612 | 0.0060963 |
| Ppp2ca | -2.2694 | 0.00055692 | Dnajb11 | 5.2032 | 0.0013598 | Sptbn1 | 3.4148 | 2.22E-06 |
| Ppp2cb | -2.6161 | 0.0032709 | Rpn1 | 3.3695 | 3.25E-06 | Sf3b2 | 2.0655 | 0.0063359 |
| Ppp5c | -3.0095 | 0.0018953 | Eif2a | 3.5382 | 0.0031617 | Sfpq | 2.0668 | 0.00039714 |
| Ppp1ca | -3.6973 | 0.0030641 | Eif3a | 1.9939 | 0.00048557 | Sod1 | 1.6557 | 0.012164 |
| Ppp1cb | -2.3087 | 0.00053768 | Eif4g1 | 2.805 | 0.0051087 | Tmsb10 | 5.4958 | 0.00020885 |
| Tf | -2.1523 | 0.00080335 | Eif4g2 | 1.9436 | 0.01205 | Tmsb4x | 3.3163 | 0.0013165 |
| Serpinb6 | -2.6652 | 0.0045587 | Ewsr1 | 1.8396 | 0.0015832 | Tor1aip1 | 2.7062 | 0.0022868 |
| Sh3bgrl3 | -1.7627 | 0.013292 | Fam107b | 2.635 | 0.00030794 | Purb | 2.7221 | 0.0051049 |
| Sh3bgrl2 | -2.2651 | 0.0013871 | Fbp1 | 2.0786 | 0.00043528 | Vcp | 3.1554 | 0.0077019 |
| Acads | -1.0703 | 0.016955 | Gnpda1 | 1.0801 | 0.0054471 | Otub1 | 5.7339 | 0.00010224 |
| Sar1a | -2.0333 | 0.013157 | Gnpda2 | 1.6016 | 0.00063796 | Ubap2l | 1.8698 | 0.0078568 |
| Abhd11 | -3.7279 | 0.008756 | Glrx2 | 3.1846 | 0.0077608 | Acadvl | 2.0584 | 0.010947 |
| Atp1a1 | -2.0952 | 0.008913 | Gpx1 | 2.1154 | 0.00068834 | Vamp7 | 4.666 | 0.015039 |
| Atp1b1 | -2.3003 | 0.0038747 | Gss | 1.6127 | 0.00015392 | Pepd | 1.5346 | 0.0018057 |
| Slco1a1 | -2.8472 | 0.0016493 | Gapdh | 2.4673 | 0.0032243 | | | |
| Sri | -4.4032 | 0.015486 | Gapdhrt | 1.9304 | 0.0023352 | | | |
| Ddx39b | -3.7166 | 0.004785 | Gapdhs | 4.2703 | 7.07E-05 | | | |
| Spryd4 | -2.5512 | 0.01627 | Gyg1 | 1.1551 | 0.0056679 | | | |
| Stard10 | -2.4633 | 0.0074394 | Gchfr | 1.1993 | 0.0028467 | | | |
| Stard4 | -5.565 | 9.48E-05 | Hras | 1.5145 | 0.00038754 | | | |
| Stard10 | -1.964 | 0.016261 | Kras | 2.5789 | 0.0077263 | | | |
| Scp2 | -3.0011 | 0.00068912 | Nras | 1.6659 | 0.0021454 | | | |
| Hspa9 | -1.8628 | 0.0054049 | Ran | 1.5654 | 0.0021651 | | | |
| Sdha | -1.5671 | 0.0023115 | Sgsh | 2.8793 | 0.005842 | | | |
| Suclg1 | -3.2377 | 0.0002632 | H1-3 | 3.0957 | 0.0073182 | | | |
| Sucla2 | -2.9354 | 0.0038661 | H1-4 | 5.8612 | 0.00012794 | | | |
| Suclg2 | -2.5952 | 0.0046244 | H2ac11 | 4.1612 | 0.00080061 | | | |
| Oxct1 | -5.3976 | 8.86E-05 | H2bc23 | 2.7341 | 0.0058333 | | | |
| Sult1c2 | -5.3373 | 0.0023392 | Adhfe1 | 1.8922 | 0.010319 | | | |
10

### Slide 11
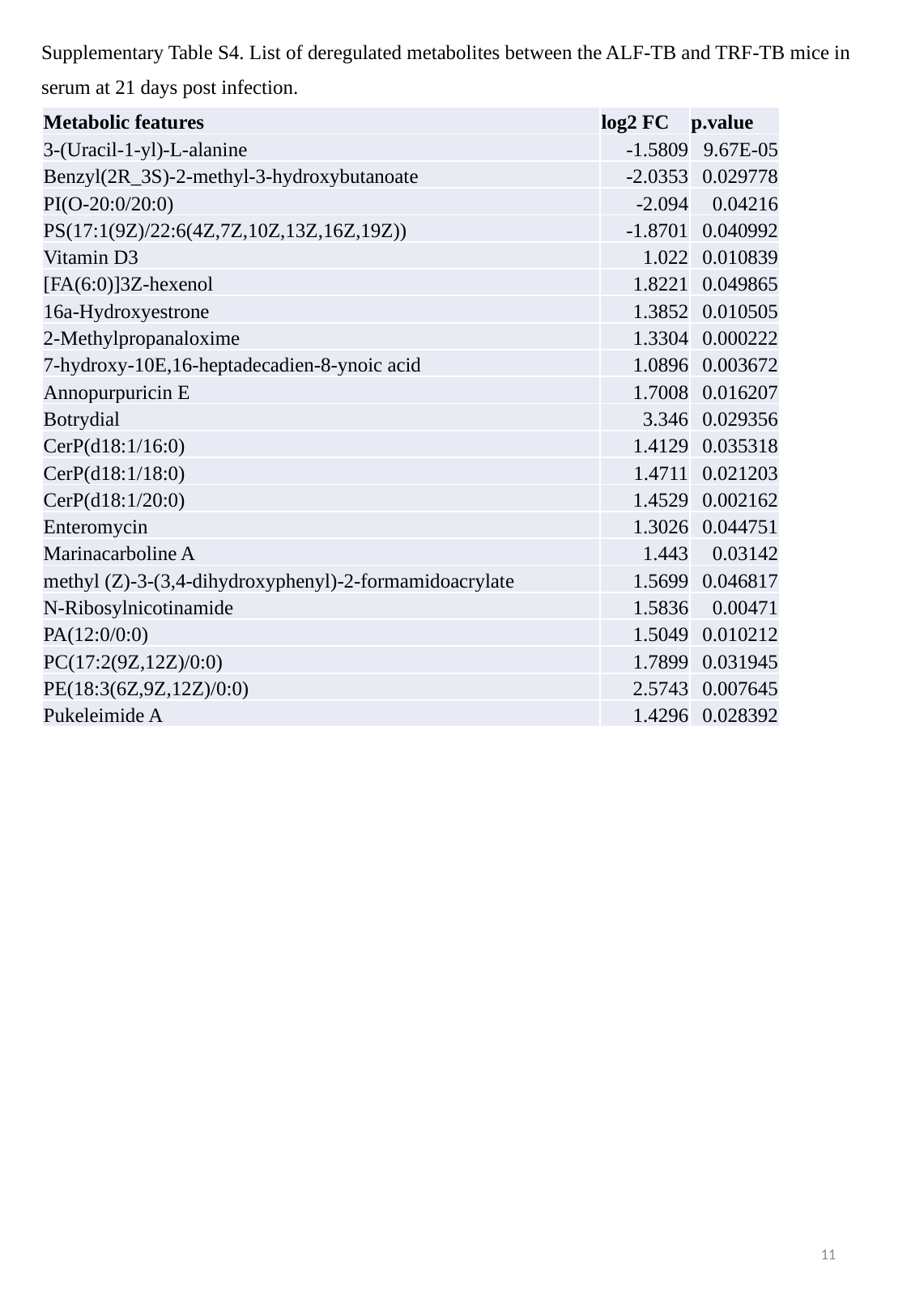

Supplementary Table S4. List of deregulated metabolites between the ALF-TB and TRF-TB mice in serum at 21 days post infection.
| Metabolic features | log2 FC | p.value |
| --- | --- | --- |
| 3-(Uracil-1-yl)-L-alanine | -1.5809 | 9.67E-05 |
| Benzyl(2R\_3S)-2-methyl-3-hydroxybutanoate | -2.0353 | 0.029778 |
| PI(O-20:0/20:0) | -2.094 | 0.04216 |
| PS(17:1(9Z)/22:6(4Z,7Z,10Z,13Z,16Z,19Z)) | -1.8701 | 0.040992 |
| Vitamin D3 | 1.022 | 0.010839 |
| [FA(6:0)]3Z-hexenol | 1.8221 | 0.049865 |
| 16a-Hydroxyestrone | 1.3852 | 0.010505 |
| 2-Methylpropanaloxime | 1.3304 | 0.000222 |
| 7-hydroxy-10E,16-heptadecadien-8-ynoic acid | 1.0896 | 0.003672 |
| Annopurpuricin E | 1.7008 | 0.016207 |
| Botrydial | 3.346 | 0.029356 |
| CerP(d18:1/16:0) | 1.4129 | 0.035318 |
| CerP(d18:1/18:0) | 1.4711 | 0.021203 |
| CerP(d18:1/20:0) | 1.4529 | 0.002162 |
| Enteromycin | 1.3026 | 0.044751 |
| Marinacarboline A | 1.443 | 0.03142 |
| methyl (Z)-3-(3,4-dihydroxyphenyl)-2-formamidoacrylate | 1.5699 | 0.046817 |
| N-Ribosylnicotinamide | 1.5836 | 0.00471 |
| PA(12:0/0:0) | 1.5049 | 0.010212 |
| PC(17:2(9Z,12Z)/0:0) | 1.7899 | 0.031945 |
| PE(18:3(6Z,9Z,12Z)/0:0) | 2.5743 | 0.007645 |
| Pukeleimide A | 1.4296 | 0.028392 |
11

### Slide 12
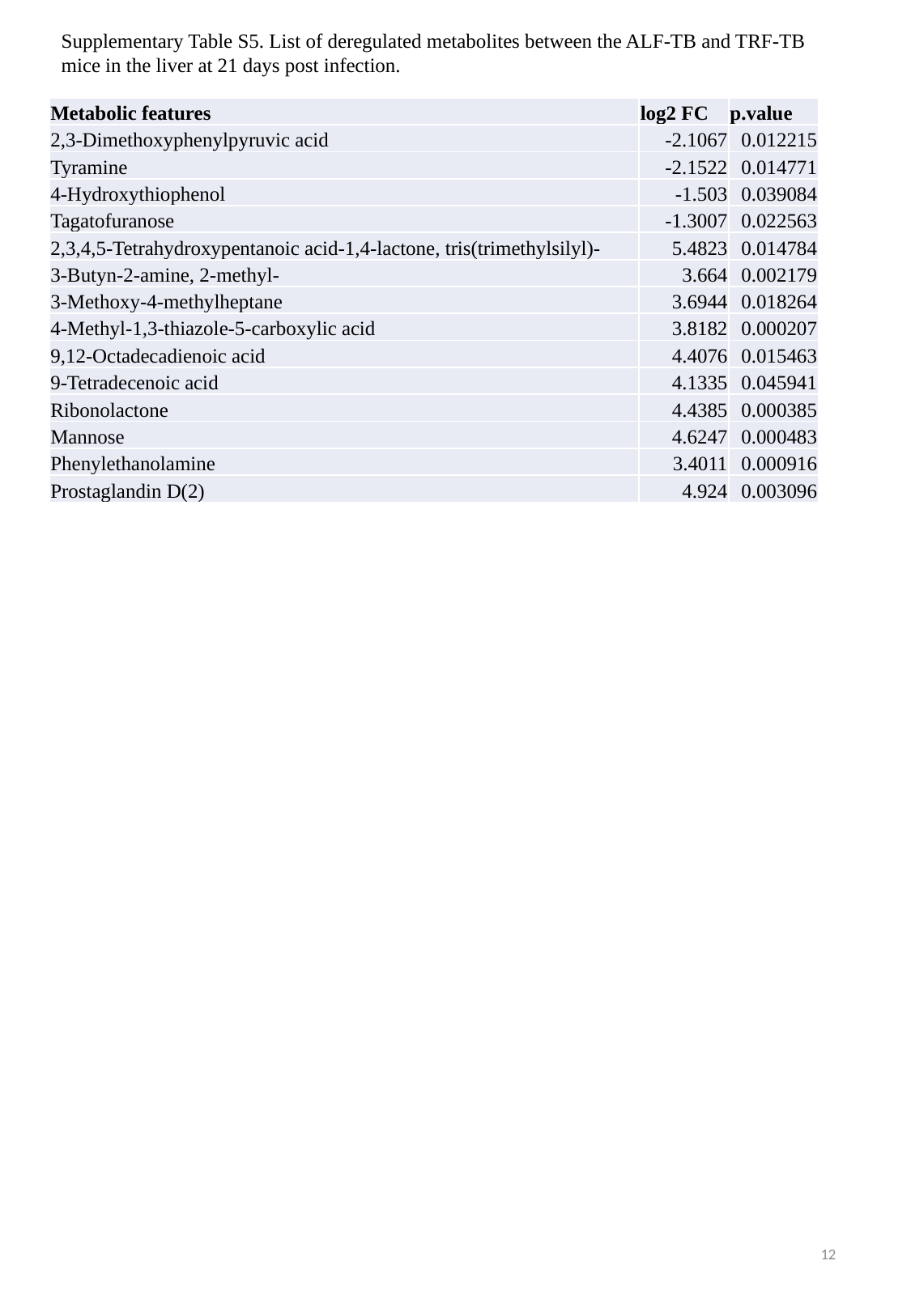

Supplementary Table S5. List of deregulated metabolites between the ALF-TB and TRF-TB mice in the liver at 21 days post infection.
| Metabolic features | log2 FC | p.value |
| --- | --- | --- |
| 2,3-Dimethoxyphenylpyruvic acid | -2.1067 | 0.012215 |
| Tyramine | -2.1522 | 0.014771 |
| 4-Hydroxythiophenol | -1.503 | 0.039084 |
| Tagatofuranose | -1.3007 | 0.022563 |
| 2,3,4,5-Tetrahydroxypentanoic acid-1,4-lactone, tris(trimethylsilyl)- | 5.4823 | 0.014784 |
| 3-Butyn-2-amine, 2-methyl- | 3.664 | 0.002179 |
| 3-Methoxy-4-methylheptane | 3.6944 | 0.018264 |
| 4-Methyl-1,3-thiazole-5-carboxylic acid | 3.8182 | 0.000207 |
| 9,12-Octadecadienoic acid | 4.4076 | 0.015463 |
| 9-Tetradecenoic acid | 4.1335 | 0.045941 |
| Ribonolactone | 4.4385 | 0.000385 |
| Mannose | 4.6247 | 0.000483 |
| Phenylethanolamine | 3.4011 | 0.000916 |
| Prostaglandin D(2) | 4.924 | 0.003096 |
12

### Slide 13
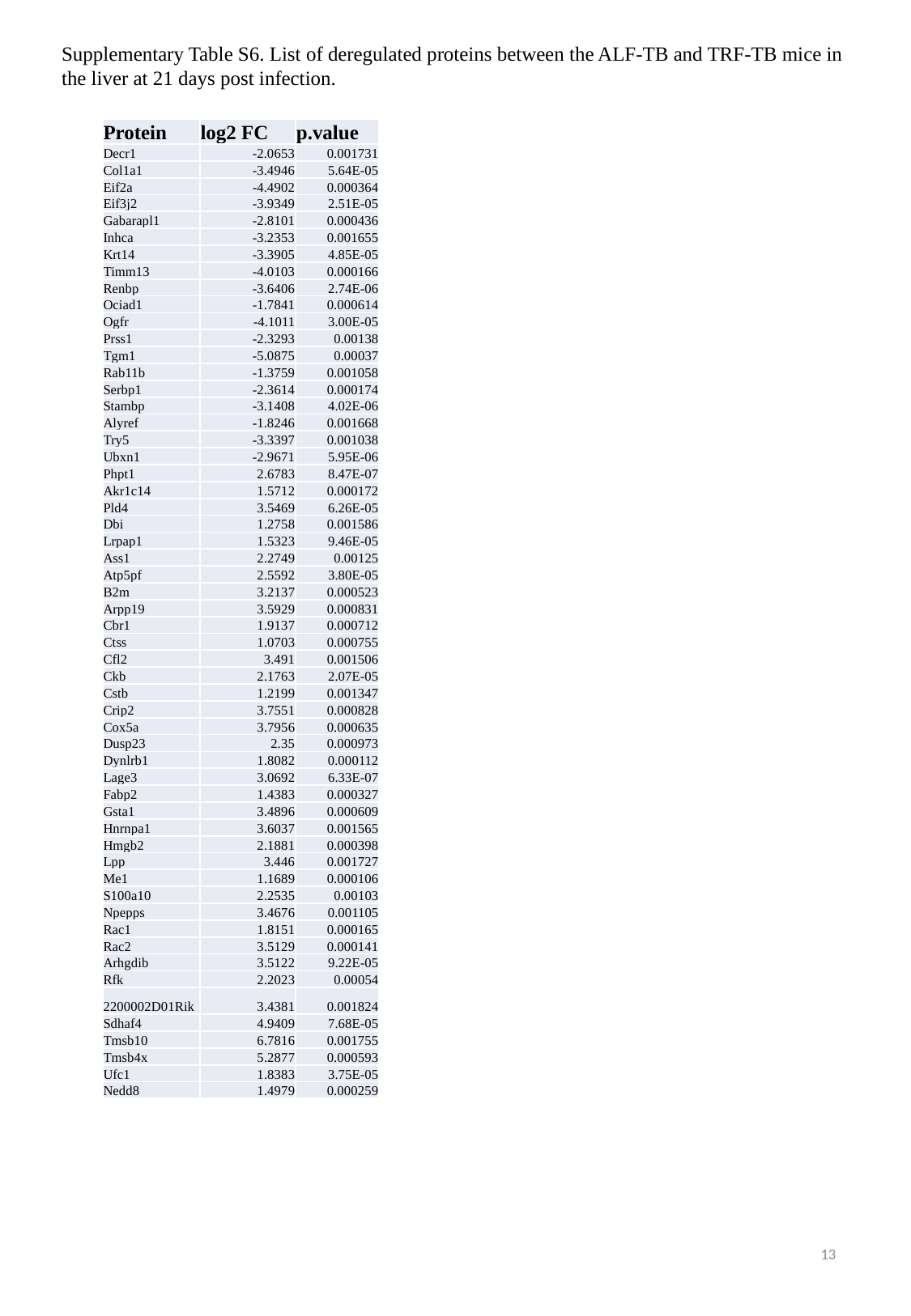

Supplementary Table S6. List of deregulated proteins between the ALF-TB and TRF-TB mice in the liver at 21 days post infection.
| Protein | log2 FC | p.value |
| --- | --- | --- |
| Decr1 | -2.0653 | 0.001731 |
| Col1a1 | -3.4946 | 5.64E-05 |
| Eif2a | -4.4902 | 0.000364 |
| Eif3j2 | -3.9349 | 2.51E-05 |
| Gabarapl1 | -2.8101 | 0.000436 |
| Inhca | -3.2353 | 0.001655 |
| Krt14 | -3.3905 | 4.85E-05 |
| Timm13 | -4.0103 | 0.000166 |
| Renbp | -3.6406 | 2.74E-06 |
| Ociad1 | -1.7841 | 0.000614 |
| Ogfr | -4.1011 | 3.00E-05 |
| Prss1 | -2.3293 | 0.00138 |
| Tgm1 | -5.0875 | 0.00037 |
| Rab11b | -1.3759 | 0.001058 |
| Serbp1 | -2.3614 | 0.000174 |
| Stambp | -3.1408 | 4.02E-06 |
| Alyref | -1.8246 | 0.001668 |
| Try5 | -3.3397 | 0.001038 |
| Ubxn1 | -2.9671 | 5.95E-06 |
| Phpt1 | 2.6783 | 8.47E-07 |
| Akr1c14 | 1.5712 | 0.000172 |
| Pld4 | 3.5469 | 6.26E-05 |
| Dbi | 1.2758 | 0.001586 |
| Lrpap1 | 1.5323 | 9.46E-05 |
| Ass1 | 2.2749 | 0.00125 |
| Atp5pf | 2.5592 | 3.80E-05 |
| B2m | 3.2137 | 0.000523 |
| Arpp19 | 3.5929 | 0.000831 |
| Cbr1 | 1.9137 | 0.000712 |
| Ctss | 1.0703 | 0.000755 |
| Cfl2 | 3.491 | 0.001506 |
| Ckb | 2.1763 | 2.07E-05 |
| Cstb | 1.2199 | 0.001347 |
| Crip2 | 3.7551 | 0.000828 |
| Cox5a | 3.7956 | 0.000635 |
| Dusp23 | 2.35 | 0.000973 |
| Dynlrb1 | 1.8082 | 0.000112 |
| Lage3 | 3.0692 | 6.33E-07 |
| Fabp2 | 1.4383 | 0.000327 |
| Gsta1 | 3.4896 | 0.000609 |
| Hnrnpa1 | 3.6037 | 0.001565 |
| Hmgb2 | 2.1881 | 0.000398 |
| Lpp | 3.446 | 0.001727 |
| Me1 | 1.1689 | 0.000106 |
| S100a10 | 2.2535 | 0.00103 |
| Npepps | 3.4676 | 0.001105 |
| Rac1 | 1.8151 | 0.000165 |
| Rac2 | 3.5129 | 0.000141 |
| Arhgdib | 3.5122 | 9.22E-05 |
| Rfk | 2.2023 | 0.00054 |
| 2200002D01Rik | 3.4381 | 0.001824 |
| Sdhaf4 | 4.9409 | 7.68E-05 |
| Tmsb10 | 6.7816 | 0.001755 |
| Tmsb4x | 5.2877 | 0.000593 |
| Ufc1 | 1.8383 | 3.75E-05 |
| Nedd8 | 1.4979 | 0.000259 |
13
